# Sustained epigenetic rejuvenation of serially engrafting human iPSC-derived HSCs

**DOI:** 10.64898/2026.07.15.732710

**Authors:** Arjun Jain, Jacky Li, Xiaojie Yu, Adeleye Opejin, Darren Yu, Anastasia Vavilina-Halstead, Alex Trapp, Jared Tumiel, Zachary Chiang, Esequiel Pastrana, Caroline Polanco, Jorge Pachas, Miguel Pulido, Bianca Carapia, Fredrick Lopez, Sanskruti Deshmukh, Mahmoud Dabbah, Alejandra Sevilla, Swathi Karthikeyan, Anastasia Shindyapina

**Author notes:** co–corresponding authors.

## Abstract

Hematopoietic stem cell (HSC) function declines with age, contributing to immunosenescence and inferior transplantation outcomes. Here, we generated iPSC-derived HSCs (iHSCs) from multiple adult donors and performed integrated epigenetic, transcriptional, telomeric, and functional analyses to see if they retain youthful identity across differentiation and serial transplantation. Longitudinal DNA methylation profiling revealed that, independent of donor age, epigenetic age was reset to near zero in iPSCs and remained under seven years across differentiation and transplantation. In contrast, hematopoietic identity was established through a two-phase process: directional remodeling during in vitro differentiation extinguished pluripotency programs and initiated hematopoietic regulatory networks, while long-term engraftment was associated with a second wave of promoter methylation differences that converged toward primary adult HSCs. Notably, methylation at age-associated sites and global entropy remained stable across both phases, and single-cell telomere analysis demonstrated restoration of telomere length in iHSCs compared to primary adult HSCs. Youthful epigenetic features were maintained through secondary transplantation. These findings demonstrate that long-term HSC identity can be achieved independently of epigenetic aging and establish a framework for evaluating rejuvenated stem cell-derived grafts in regenerative medicine.

## Introduction

Hematopoietic stem cell (HSC) transplantation is curative for malignant and non-malignant blood disorders, but its broader application is constrained by donor availability, graft-versus-host disease, and donor-dependent variation in stem cell function [Copelan, 2006; Gratwohl et al, 2015; Snowden et al, 2022]. Most clinical grafts are derived from mobilized peripheral blood or bone marrow of adult donors, and donor age is a major determinant of transplant outcome [Kollman et al, 2016; DeZern et al, 2021; Shaw et al, 2018; Abid et al, 2023]. At the cellular level, hematopoietic aging is associated with reduced HSC regenerative capacity, myeloid-biased differentiation, impaired lymphoid output, accumulated DNA damage, and altered clonal composition [Rossi et al, 2005; Pang et al, 2011; Kuranda et al, 2011; Rübe et al, 2011; Flach et al, 2014].

These changes are accompanied by alterations in DNA methylation, including gains at Polycomb-regulated loci and increased epigenetic entropy [Beerman et al, 2013; Sun et al, 2014]. DNA methylation-based clocks allow us to accurately measure chronological and biological aging in blood and hematopoietic tissues [Horvath, 2013; Hannum et al, 2013; Levine et al, 2018]. After allogeneic transplantation, epigenetic age measurements in recipient blood reflect donor–intrinsic properties and can show transient acceleration or remodeling during immune reconstitution [Weider et al, 2014; Stölzel et al, 2017; Søraas et al, 2019].

In addition to DNA methylation changes, telomere length declines with age in human hematopoietic cells [Vaziri et al, 1993; Vaziri et al, 1994; Aubert et al, 2012], and inherited telomere biology disorders result in bone marrow failure [Yamaguchi et al, 2005]. In the transplantation setting, donor telomere length has been associated with post-transplant survival, directly linking telomere reserve to hematopoietic graft performance [Gadalla et al, 2015; Gadalla et al, 2018; Gadalla et al, 2025].

Reprogramming of somatic cells into induced pluripotent stem cells (iPSCs) resets epigenetic state and extends telomeres, erasing age-associated DNA methylation signatures and reactivating telomerase. As a result, iPSCs exhibit near-zero epigenetic ages and embryonic-like telomeres [Takahashi et al, 2007; Horvath, 2013; Marion et al, 2009; Suhr et al, 2009]. However, whether iPSC-derived HSCs can acquire definitive long-term HSC identity while maintaining young-like epigenetic and telomere features remains unexplored. Definitive HSCs arise during development from arterial hemogenic endothelium and acquire long-term repopulating capacity through niche-dependent maturation [Ivanovs et al, 2011; Ditadi et al, 2015; Zeng et al, 2019], and recent single-cell atlases have defined conserved transcriptional trajectories that distinguish nascent, fetal, and adult HSC states [Calvanese et al, 2022].

A recent study described differentiation conditions that generate transgene-free iHSCs capable of long-term multilineage engraftment in immunodeficient mice [Ng et al, 2024]. While this work established functional engraftment of iHSCs, how epigenetic age and HSC identity are remodeled during differentiation and subsequent in vivo maturation, and whether embryonic epigenetic features are preserved under proliferative and transplantation stress, remains unresolved.

Here, we generate iHSCs from multiple adult-derived human iPSC lines and characterize epigenetic, transcriptional, and functional state of cells throughout differentiation and primary and secondary transplantation. By combining longitudinal DNA methylation profiling with single-cell RNA sequencing, telomere length measurements, and in vivo functional assays, we show that hematopoietic identity is acquired through a two-phase process – initiation during in vitro differentiation and completion following exposure to the bone marrow niche – while preserving epigenetic age under seven years irrespective of donor age. These findings demonstrate that iPSC-derived HSCs preserve epigenetic and telomeric characteristics of iPSCs through downstream progeny and serial transplantation, indicating that functional and rejuvenated hematopoietic grafts can be generated across donors.

## Results

### iPSC-derived HSCs from adult donors remain epigenetically young across differentiation and engraftment

To characterize the epigenetic changes in human iHSCs across differentiation and engraftment, we obtained iPSC lines from four young (ages 18-26) and two old (ages 57-60) donors and differentiated them into iHSCs using a swirling embryoid body protocol optimized for engraftment [Ng et al, 2024] (Fig. 1A, left). We transplanted resulting iHSCs from all donors retro-orbitally into >5 female NOD.Cg-*Kit^W-41J^ Tyr^+^ Prkdc^scid^ Il2rg^tm1Wjl^*/ThomJ (NBSGW) mice each (5 million cells/mouse), which support engraftment of human HSCs without irradiation [McIntosh et al, 2015]. Mice were followed for 20 weeks, with routine bleeds at 12 and 16 weeks post-transplant to track human hematopoietic chimerism, and week 20 terminal bone marrow, spleen, and peripheral blood were analyzed for human engraftment. (Fig. 1A, right). iHSCs from each donor were profiled across differentiation and transplant via flow cytometry, and scRNA-sequencing was performed for all iHSC lots. DNA methylation profiling was conducted for three donors at various time points across differentiation, and on iHSCs from all donors after long-term engraftment. Finally, iHSCs from three donors were also subject to single-cell telomere analysis.

**FIGURE 1.**
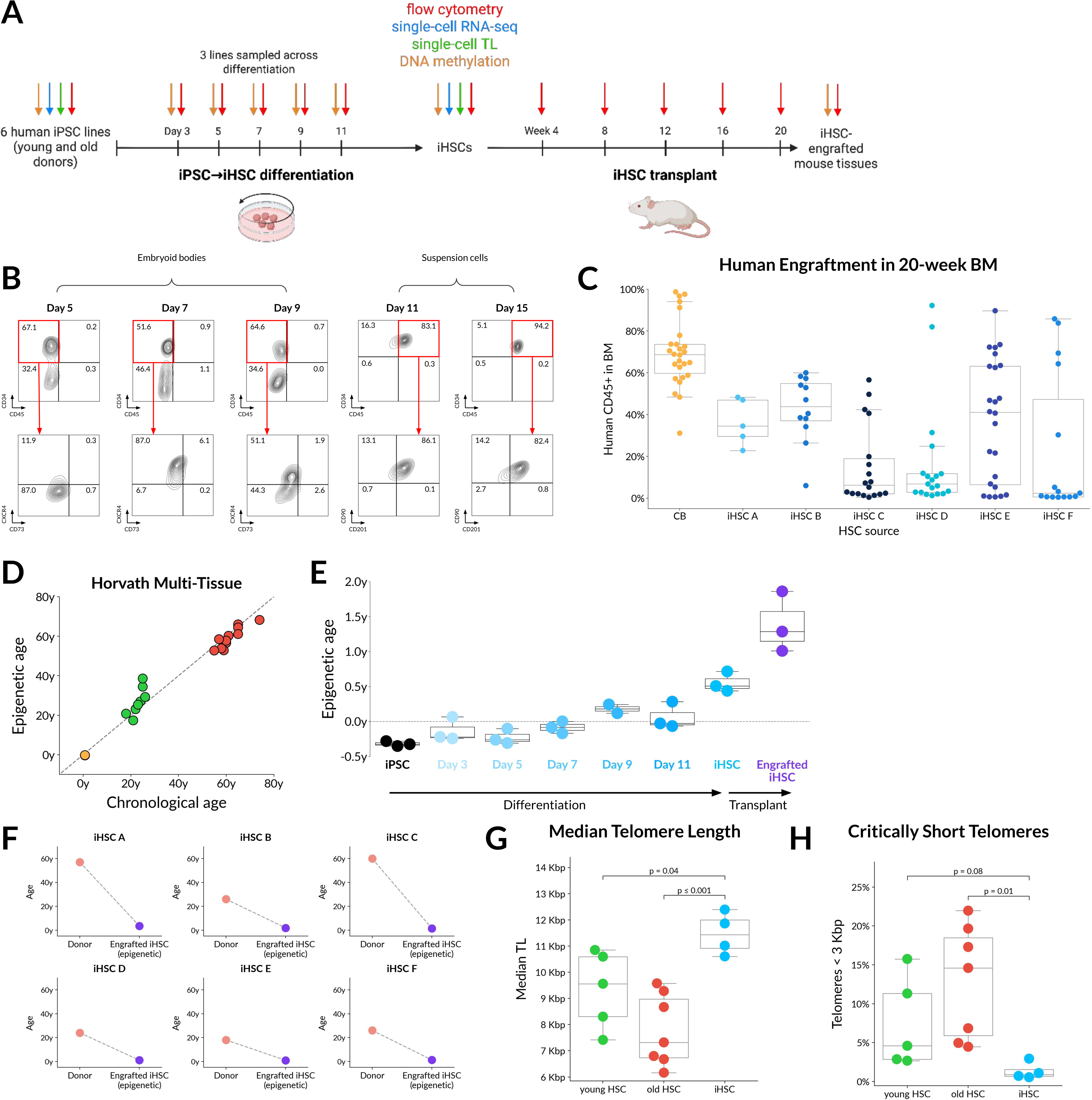
**(A)** Experimental workflow. Human iPSC lines derived from adult donors were differentiated into iHSCs using a swirling embryoid body protocol and transplanted into NBSGW mice for 20-week engraftment. Samples were collected at labeled timepoints for flow cytometry, DNA methylation profiling, single-cell RNA-sequencing, and single-cell telomere length analysis. **(B)** Flow cytometric characterization of iPSC to iHSC differentiation (representative differentiation of iPSC A shown). Early CD34+ hemogenic endothelial populations emerge in embryoid bodies, followed by appearance of CD34+CD45+ hematopoietic cells in suspension by day 11. Day 15 iHSCs express HSC-associated markers including CD90 and CD201/EPCR. **(C)** Human CD45+ engraftment in NBSGW bone marrow at 20 weeks post-transplant. Each dot represents one mouse. **(D)** Horvath multi-tissue epigenetic clock predictions across fetal (orange), young (green), and old (red) primary CD34+ HSCs demonstrating high correlation with chronological age. **(E)** Horvath-predicted epigenetic age during differentiation of three iPSC lines. iPSCs reset to ∼0 years and gradually increase during differentiation; day 15 iHSCs and week 20 engrafted cells maintain fetal-like predicted age. **(F)** Donor age vs Horvath-predicted epigenetic age after primary engraftment. Human CD45+ cells isolated from engrafted bone marrow retain young predicted age independent of donor age. **(G)** Median telomere length (MTL) of day 15 iHSCs versus young and old primary HSCs measured by single-cell HT Q-FISH. **(H)** Proportion of critically short telomeres (<3 kb) in day 15 iHSCs versus young and old primary HSCs.

For the differentiation, flow cytometry results for all donor iPSC lines matched the known trajectory [Ng et al, 2024], including the appearance of CD34+CD45+ hematopoietic cells in the suspension fraction by day 11 (Fig. 1B). Single-cell RNA-sequencing of day 15 cells confirmed expression of signature HSC transcription factors RUNX1, HOXA9, MLLT3, MECOM, HLF, and SPINK2 [Calvanese et al, 2022] (Fig. S1A). 94 of 106 NBSGW mice transplanted with iHSCs had >1% human engraftment in week 20 bone marrow (average 26.37 ± 25.93%), and analysis of week 20 mouse tissues demonstrated robust multilineage reconstitution, including lymphoid (CD19+, CD3+), myeloid (CD33+), and erythroid (CD235a+) compartments, at levels comparable to those observed in transplantation of cord blood- (CB) derived human CD34+ cells (Fig. 1C, S1B). The bone marrow of iHSC-engrafted mice displayed hematopoietic progenitor lineages, including a CD45+CD34+CD38– HSPC compartment (Fig. S1C). While engraftment levels varied across mice, this variability was consistent with xenotransplants of adult primary HSCs (Fig. S1D) and previously published results [Ng et al, 2024].

We then asked how epigenetic age changed across this process. DNA methylation-based age estimates were calculated with three PC-based epigenetic clocks [Higgins-Chen et al, 2022], including the Horvath pan-tissue clock [Horvath, 2013], which accurately predicted the chronological age of our fetal and adult primary HSC samples (r = 0.98) with an average error of 3.46 ± 3.16 years (Fig. 1D). As previously reported [Horvath, 2013], the epigenetic age of iPSCs was predicted to be near zero (−0.32 ± 0.04 years), and it gradually increased during differentiation (Fig. 1E). The iHSCs harvested at day 15 of differentiation maintained a young epigenetic age, predicted as 0.55 ± 0.14 years old by the Horvath clock (Fig. 1E).

Notably, after transplant, engrafted iHSC-derived blood maintained a young epigenetic age. The average epigenetic age of human CD45+ cells isolated from engrafted mouse bone marrow was 1.68 ± 0.93 years (Fig. 1F). For the three donors profiled across the entire workflow, this represented an increase of 0.29-1.42 years across transplantation, or 2.17 ± 1.48 days per day in vivo on average (Fig. S1F). This is in line with our results from primary CB-derived CD34+ cells, whose epigenetic age increased at the rate of 2.11 days per day in vivo across long-term engraftment (Fig. S1F), as well as with published Horvath clock results after human cord blood transplants [Onizuka et al, 2023]. The Hannum [Hannum et al, 2013] and PhenoAge [Levine et al, 2018], epigenetic clocks similarly predicted young ages for iHSCs and their engrafted progeny (Fig. S1E, S1G).

Finally, we sought to measure the telomere length of iPSC-derived HSCs after differentiation, as it has been previously shown that the process of reprogramming somatic cells into iPSCs reactivates telomerase and extends telomeres [Marion et al, 2009; Suhr et al, 2009]. The telomeres of day 15 iHSCs were characterized at a single-cell level via high-throughput quantitative fluorescence in situ hybridization (HT Q-FISH) [Canela et al, 2007; De Pedro et al, 2020] alongside adult primary human HSCs. iHSCs maintained a significantly longer median telomere length (p = 0.0022, two-sided Wilcoxon rank-sum test) compared to primary HSCs (11,469 bp and 8,435 bp, respectively, Fig. 1G), as part of a shift across the entire telomere length distribution (Fig. S1I). In particular, the proportion of critically short telomeres (<3 kb) was significantly lower in iHSCs (p = 0.0044, two-sided Wilcoxon rank-sum test), with just 1.33% of telomeres on average under this threshold versus 10.59% in adult primary HSCs (Fig. 1H).

Collectively, these findings indicate that functional iPSC-derived HSCs from adult donors maintain youthful molecular and epigenetic features across differentiation and engraftment, despite the burden of transplant and reconstitution in vivo.

### iHSC identity convergence in vivo is decoupled from epigenetic aging

We next explored how hematopoietic identity is established during differentiation and whether iHSC maturation is completed in vitro or requires in vivo niche exposure. Although iHSCs maintain an epigenetic age under 7 years old through differentiation and transplantation, we observed global remodeling of DNA methylation associated with hematopoietic identity.

Principal component analysis (PCA) of genome-wide DNA methylation profiles revealed that in vitro differentiation from iPSCs to iHSCs induced a directional shift along a major identity-associated axis (PC3), moving cells away from pluripotency and toward a hematopoietic state (Fig. 2A). However, in vitro-derived iHSCs remained epigenetically distinct from primary human HSCs, clearly seen across PC1 (Fig. 2A). Following long-term engraftment in immunodeficient mice, iHSC-derived cells underwent a pronounced additional shift in DNA methylation space, converging with young and old primary human HSCs along both identity-associated PCs (Fig. 2A). This engraftment-associated convergence was directional and distinct from the in vitro differentiation trajectory, consistent with acquisition or enrichment of a primary-HSC-like epigenetic state after transplantation.

**FIGURE 2.**
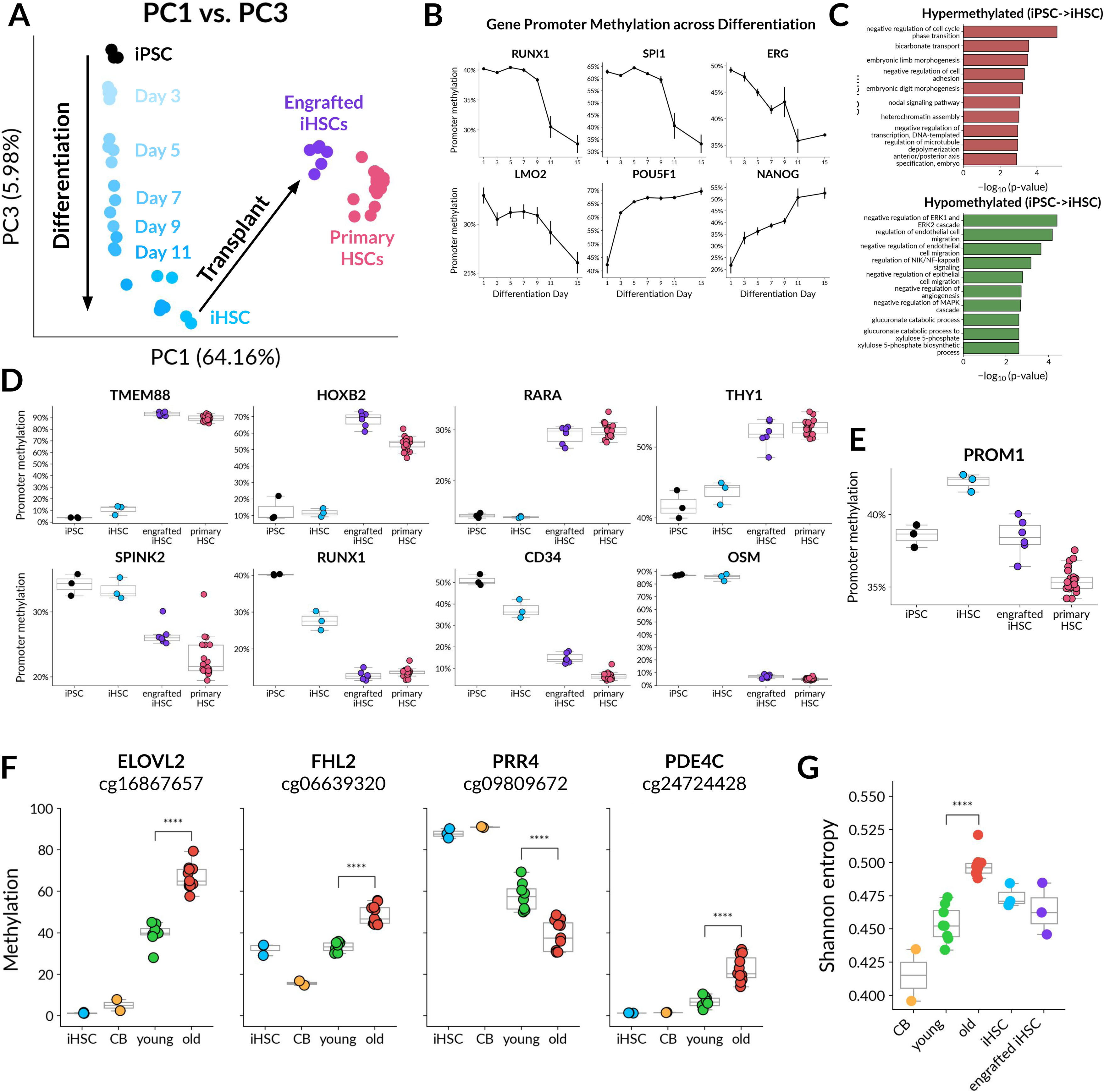
**(A)** Principal component analysis of genome-wide DNA methylation profiles. In vitro differentiation drives directional movement along PC3 toward hematopoietic identity while engraftment induces additional convergence toward primary HSCs along PC1 and PC3. **(B)** Examples of mean promoter methylation dynamics across differentiation for representative pluripotency and hematopoietic genes. **(C)** Gene ontology enrichment of promoters hypermethylated (top) or hypomethylated (bottom) during iHSC differentiation. **(D)** Promoters remodeled primarily during engraftment, including canonical HSC regulators. Methylation trajectories demonstrate incomplete remodeling in vitro and convergence after niche exposure. **(E)** Biphasic remodeling example of PROM1 (CD133) promoter, showing hypermethylation during differentiation and hypomethylation following engraftment. **(F)** Age-associated CpG loci (ELOVL2, FHL2, PRR4, PDE4C) remain stable and youthful across differentiation and engraftment. **(G)** Methylation entropy analysis demonstrating lower epigenetic entropy in iHSCs compared to aged primary HSCs across all stages.

Promoter-level methylation analysis further resolved these dynamics. By evaluating differentially methylated gene promoters and grouping CpG sites according to their methylation trajectories across differentiation and transplant, we identified three principal classes of behavior.

The first group consisted of gene promoters that were epigenetically remodeled during in vitro differentiation. From the initial iPSCs to day 15 iHSCs, 585 gene promoters were differentially methylated (|LFC|>1, FDR<0.05), as pluripotency-associated genes (such as POU5F1 and NANOG) were hypermethylated early and canonical hematopoietic regulators known to operate during the endothelial-to-hematopoietic transition and early HSC establishment, including RUNX1, SPI1, ERG, and LMO2, were hypomethylated later (Fig. 2B). Gene ontology analysis of hypermethylated promoters across differentiation supported this transition, as these loci were significantly enriched for biological processes related to embryonic development, stem cell maintenance, and early patterning programs (Fig. 2C, up), consistent with epigenetic repression of pluripotency and non-hematopoietic developmental trajectories as cells commit to the hematopoietic lineage. Conversely, promoters that became hypomethylated during differentiation were enriched for processes associated with hemogenic endothelial development and differentiation (Fig. 2C, down). Together these results indicate that in vitro differentiation induced large, directional epigenetic changes consistent with the silencing of pluripotency-associated programs and the establishment of core hematopoietic regulatory features. However, despite this substantial identity remodeling, in vitro-derived iHSCs remained epigenetically distinct from primary human HSCs, as more than 8,866 gene promoters were differentially methylated between the two groups, which is reflected by the PCA (Fig. 2A).

A second class of loci differed following long-term engraftment. Promoters of 5,902 genes in engrafted iHSC-derived cells significantly differed from pre-transplant iHSCs after 20 weeks, and 91% of these differences were directionally toward the epigenetic state of primary HSCs (Fig. S2A). Genes in this category included multiple regulators associated with definitive HSC identity and long-term hematopoietic maintenance, such as TMEM88, HOXB2, RARA, THY1 (CD90), SPINK2, RUNX1, CD34, and OSM (Fig 2D). Methylation at several of these promoters changed across differentiation but did not reach the level of primary HSCs until after engraftment (Fig. 2D). This extended into immune signaling and lineage-identity genes, including strong changes at CST7, CNR2, CD6, LAT2, and LILRA1. Accordingly, gene ontology analysis of differentially methylated promoters following engraftment revealed enrichment for biological processes related to immune and cytokine signaling and downstream blood cell function, reflecting engagement of physiological signaling pathways encountered during in vivo hematopoiesis but absent from the in vitro differentiation environment (Fig. S2B).

A subset of loci in this group showed opposite changes across these two phases, such as PROM1 (CD133), which was hypermethylated across differentiation but hypomethylated during transplant (Fig. 2E). Although PROM1 is expressed in subsets of human pluripotent stem cells and early progenitors, it is frequently downregulated during early differentiation [Vodyanik et al, 2005] and is lowly expressed in day 15 iHSCs relative to primary HSCs. However, PROM1 is a canonical marker for mature human HSCs, enriching for repopulating cells [Yin et al, 1997; Gorgens et al, 2013; Cimato et al, 2018; Calvanese et al, 2022]. The biphasic promoter methylation pattern observed here is therefore consistent with transient repression of PROM1 during progenitor stages followed by restoration or enrichment of a more functional HSC regulatory state after engraftment. This kind of two-step behavior supports the broader conclusion that the engrafted graft contains regulatory states more closely aligned with primary HSCs than the pre-transplant iHSC product.

Finally, in contrast to promoters undergoing differentiation- or engraftment-associated remodeling, we identified a third class of CpG sites whose methylation state remained stable across both in vitro differentiation and in vivo transplantation, despite extensive changes in hematopoietic identity. This group was strongly enriched for age-associated CpG sites previously shown to track chronological or biological aging across tissues, as suggested by the young epigenetic clock predictions (Fig. 1F, S1E, S1G). Canonical age-informative loci, including CpG sites within ELOVL2, FHL2, PRR4, and PDE4C [Horvath, 2013; Hannum et al, 2013; Garagnani et al, 2012; Zbieć-Piekarska et al, 2015; Weidner et al, 2014; Bocklandt et al, 2011], retained youthful methylation states in iHSCs relative to old primary HSCs and did not acquire the age-associated methylation gains observed in adult hematopoietic stem cells, even after long-term engraftment (Fig. 2F). Consistent with this stability, measures of methylation entropy [Hannum et al, 2013], which increases with age and reflects stochastic drift and loss of epigenomic integrity in primary HSCs (Fig. 2G), remained significantly reduced in iHSCs throughout differentiation and following transplantation compared with aged primary HSCs (Fig. 2G).

In addition to chronological epigenetic clocks, we examined methylation-derived measures of aging dynamics, including DunedinPACE, a blood-trained rate-of-aging estimator derived from longitudinal biomarker change [Belsky et al, 2022], and SystemsAge, a composite epigenetic aging score derived from blood methylation profiles [Sehgal et al, 2025]. DunedinPACE was elevated in iPSCs and day 15 iHSCs relative to primary hematopoietic samples (Fig. S2C). However, following engraftment, DunedinPACE values decreased and converged toward those observed in primary young and old mPB HSCs (Fig. S2C). Similarly, elevated SystemsAge scores were dramatically reduced after 20 weeks in vivo compared to in vitro states (Fig. S2D). While chronological epigenetic age remained low throughout differentiation and engraftment, pace-of-aging signatures normalized upon niche exposure.

Thus, iPSC-derived HSC grafts show separable changes in hematopoietic identity and epigenetic aging. In vitro differentiation initiates directional remodeling that extinguishes pluripotency programs but remains insufficient to fully recapitulate the definitive HSC epigenetic state. After transplantation, the engrafted human graft converges toward primary HSC methylation states, likely reflecting some combination of niche-associated maturation and selective expansion of cells with greater repopulating potential. In contrast, age-associated DNA methylation features remain stable across both differentiation and engraftment, preserving a youthful epigenetic state.

### iHSCs maintain young epigenomes through serial transplant

Finally, we evaluated the durability of iHSC function and epigenetic state through serial transplantation. Up to 10 × 10^6^ bone marrow cells of primary transplant mice engrafted with iHSCs from three donors (ages 57, 26, and 60) or CB HSC controls were collected and injected into new (secondary) NBSGW recipients (Fig. 3A) which were monitored for 20 weeks. The peripheral blood of secondary recipients showed increasing human CD45+ chimerism across 12 to 20 weeks post-transplant, up to 84% of lysed blood in an individual mouse (Fig. 3B), and bone marrow analysis at the 20-week time point revealed robust reconstitution (Fig. 3D). 18 of 23 mice showed >1% human cell engraftment in the bone marrow, ranging from 0.3-84.9% (Fig. 3D). The bone marrow, peripheral blood, and spleens contained clear erythroid (CD235a+), myeloid (CD33+), B cell (CD19+), and T/NK cell (CD3+) compartments (Fig. 3C,E). Lineage distributions were comparable to that of CB secondary transplant, with an increased proportion of CD235a+ cells (Fig. 3E). Within the human graft, a phenotypic CD45+CD34+CD38–CD90+CD133+ HSPC compartment was readily detected, consistent with long-term stem and progenitor maintenance (Fig. 3F).

**FIGURE 3.**
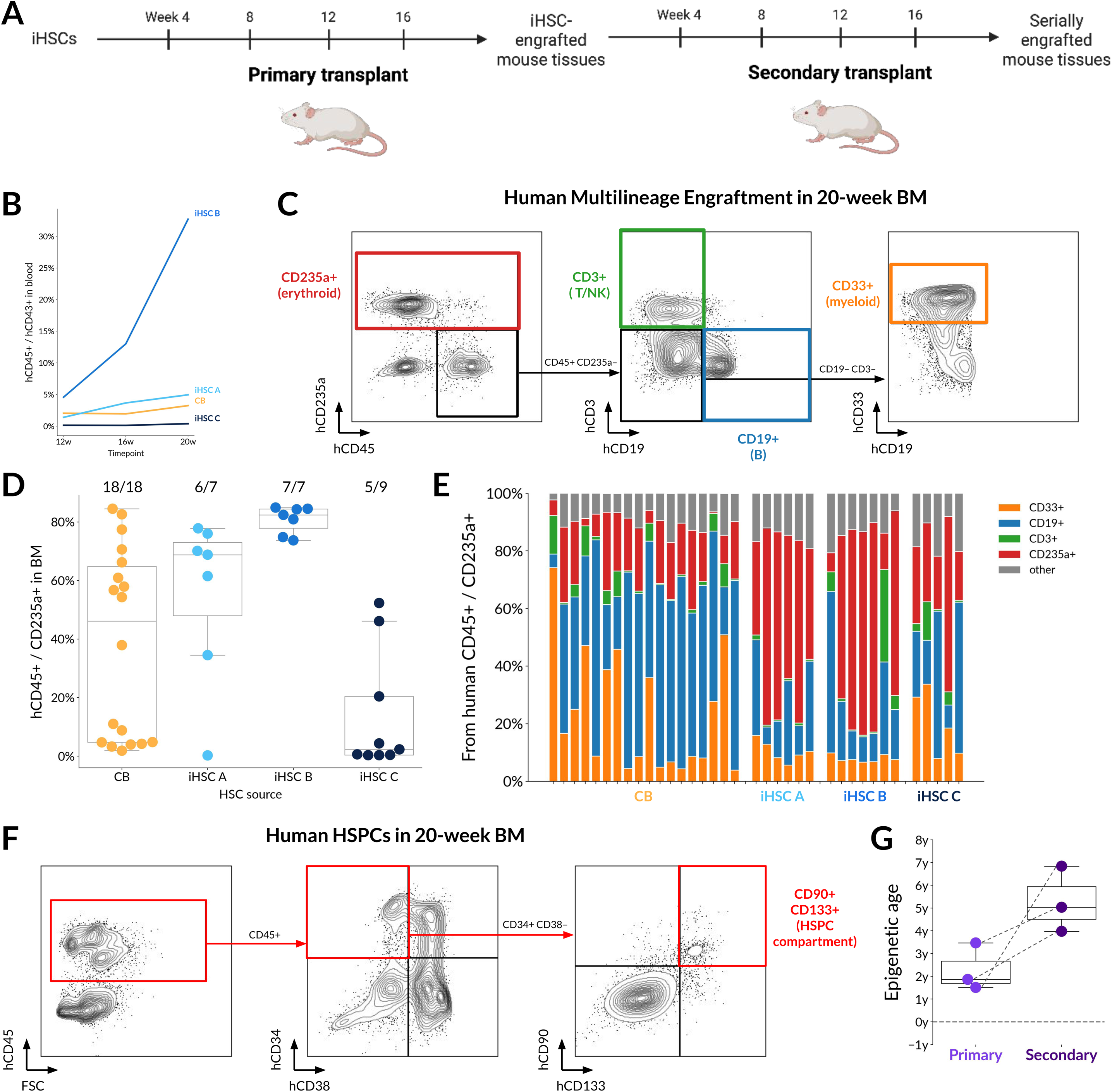
**(A)** Serial transplantation workflow. Primary engrafted bone marrow transplanted into secondary NBSGW recipients for another 20 week engraftment period. **(B)** Human CD45+ chimerism in peripheral blood of secondary recipients from 16 to 20 weeks post-transplant. **(C–E)** Multilineage reconstitution in secondary recipients across bone marrow, peripheral blood, and spleen. **(F)** Detection of CD45+CD34+CD38-CD90+CD133+ HSPC compartment in secondary bone marrow. **(G)** Horvath clock epigenetic age predictions of secondary engrafted human CD45+ cells demonstrating modest increases without reversion to donor age.

Human CD45+ iHSC-derived cells from 20-week secondary recipient bone marrow were isolated for DNA methylation profiling to determine the effects of serial proliferative stress on epigenetic age and identity. Serially engrafted iHSC-derived blood cells demonstrated retention of a young epigenetic age while identity-associated methylation patterns further aligned with primary human HSCs. Human hematopoietic cells in the bone marrow of secondary recipients had a Horvath clock-predicted age of 5.28 ± 1.45 years, which represented an increase of 3.00 ± 2.03 years during the secondary transplant on average across three donors (Fig. 3G), or 7.83 ± 5.30 days per day in vivo (Fig. S3A). For comparison, the CB graft aged 1.37 days per day in vivo. The Hannum and PhenoAge clocks predicted similar small increases in epigenetic age across secondary transplant without a reset to donor age (Fig. S3B).

Compared to the global remodeling across primary transplant, there were significantly fewer changes across secondary transplant, with only 1,543 differentially methylated promoters. The integrated PCA reflected this as well, with serially transplanted iHSC-derived cells clustering close to primary transplanted iHSC-derived cells and primary HSCs (Fig. S3C).

Together we show that iHSCs retain long-term HSC function and youthful epigenetic features despite the proliferative and selection pressures imposed by serial transplantation, supporting durable self-renewal without acquisition of age-associated methylation patterns.

## Discussion

A persistent obstacle in regenerative hematology has been reconciling three properties in the same human graft: definitive HSC identity, durable self-renewal under serial stress, and youthful molecular features that are ordinarily lost with aging. Building on the transgene-free iHSC differentiation framework of Ng and colleagues [Ng et al, 2024], our study provides evidence that polyclonal, donor-diverse iPSC-derived HSCs – including from older donors (60 years) – can generate robust multilineage engraftment, support strong secondary transplantation, and retain young-like epigenetic features across differentiation and serial transplant.

Most demonstrations of stem cell-derived hematopoiesis have relied on a small number of donor lines. By testing polyclonal iPSC lines from multiple donor ages and somatic sources (PBMCs, fibroblasts, MSCs), we reduce the likelihood that the observed phenotype is donor- or cell-type-specific. This is particularly relevant for clinical translation, where manufacturing may involve different somatic cell sources and where donor-to-donor variability is unavoidable. While our results are consistent with literature showing that youth metrics are largely independent of donor age after reprogramming to the iPSC state [Horvath, 2013; Marion et al, 2009; Suhr et al, 2009], we demonstrate that this trend extends to the long-term HSC graft level, despite stringent in vivo selection.

Secondary transplantation remains the most stringent functional readout of long-term HSC self-renewal and has historically been a bottleneck in iPSC-derived hematopoiesis studies. In the Ng et al study, secondary engraftment was limited, potentially constrained by NBSGW niche conditions and the quantity of marrow transferred to secondary recipients, as just 0.3-2.0 × 10^6^ bone marrow cells were transplanted [Ng et al, 2024; Hess et al, 2020]. With high primary engraftment and high secondary doses, up to 10 × 10^6^ cells per mouse, we show durable secondary reconstitution from multiple donors in the NBSGW model. Importantly, serially engrafted cells maintained a CD45+CD34+CD38-CD90+CD133+ compartment and multilineage output, indicating that the engrafting cells are not transient progenitors but represent self-renewing HSC activity. Secondary recipients were followed for the same 20-week duration as primary recipients, representing nearly ten months of continuous in vivo proliferation from the original iPSC-derived graft. The combination of robust multilineage reconstitution and maintenance of a stem cell immunophenotype through serial transplantation constitutes some of the strongest long-term self-renewal evidence reported to date for a transgene-free human iPSC-derived HSC platform.

Serial transplantation also provides a test of epigenetic stability. CB and iHSC grafts showed accelerated (2.1-2.2x) epigenetic aging across primary transplant, in line with the 2.7x age acceleration in the Horvath clock previously reported after human cord blood transplants [Onizuka et al, 2023]. However, the rate of epigenetic aging across secondary transplants increased for iHSC grafts relative to cord blood. The larger Horvath-age increase observed in iHSC secondary recipients relative to cord blood may reflect differences in effective LT-HSC dose, lineage composition of the bulk CD45+ compartment, or selection bottlenecks under serial stress.

Developmental studies have long emphasized that HSC competence emerges through niche interactions rather than cell-autonomous fate specification alone [Ivanovs et al, 2011; Calvanese et al, 2022], and the methylation analyses suggest that HSC identity is not fully captured by the pre-transplant iHSC product. In vitro differentiation silences pluripotency programs and activates early hemogenic and HSC regulatory modules. However, day 15 iHSCs remained epigenetically distinct from primary adult HSCs, despite expressing canonical HSC transcriptional modules defined in developmental atlases [Calvanese et al, 2022]. After engraftment, the human graft showed a second wave of promoter differences that converged toward primary HSC state, including broad changes in immune/cytokine and niche-responsive pathways. Because these measurements were performed on bulk engrafted human CD45+ cells, they cannot distinguish whether this convergence reflects cell-intrinsic maturation after niche exposure, selective expansion of a pre-existing HSC-like subset, or both. Nevertheless, both interpretations are relevant for graft engineering: the pre-transplant product contains cells capable of generating a primary-HSC-like, young epigenetic graft after long-term engraftment, and the pathways associated with this convergence remain tractable targets for improving the in vitro differentiation.

A striking result was the stability of age-associated CpGs and the maintenance of low methylation entropy across differentiation, primary transplant, and even serial transplantation. In murine systems, proliferative stress and inflammatory cues can accelerate age-like methylome remodeling in HSCs [Beerman et al, 2013; Sun et al, 2014; Yanai et al, 2024]. The persistence of young-like signatures in our iHSC grafts suggests that reprogramming establishes an epigenetic starting state that is comparatively resistant to this drift, even under the proliferative demand of engraftment.

Multiple iPSC-derived cell types, including neurons [Steg et al, 2021; Imm et al, 2021; Mendez et al, 2023], MSCs [Frobel et al, 2014], retinal organoids [Hoshino et al, 2019], and lung epithelial cells [Kabacik et al, 2022], have been shown to retain a fetal-like epigenetic age, often displaying re-initiation of epigenetic aging across in vitro differentiation. In those contexts, reprogramming resets molecular age, but adult-like maturation does not typically require extensive proliferative stress in vivo, whereas engrafted HSCs undergo sustained proliferation to reconstitute the immune system. The maintenance of low epigenetic age after ten months of primary and secondary transplantation therefore represents a more stringent test of molecular stability than most iPSC-derived differentiated systems. The present work provides evidence that a rejuvenated epigenetic state can persist in a highly proliferative adult stem cell context.

Telomere analyses reinforce this conclusion. While telomerase reactivation and telomere elongation in iPSCs have been previously documented [Marion et al, 2009; Suhr et al, 2009], the key finding here is the substantial reduction in critically short telomeres, which disproportionately drive DNA damage signaling and stem cell dysfunction [Hemann et al, 2001]. Given the clinical relevance of telomere reserve in marrow failure and transplant settings, this distribution shift suggests that iHSC grafts may possess enhanced reconstitution potential relative to adult donors.

Several limitations define next steps. Identity convergence in this study occurs within a murine niche, and evaluating alternative recipient strains or humanized niche models will clarify the extent to which the resulting state is environment-specific. Bulk CD45+ methylation profiling may partially reflect lineage composition and cannot distinguish in vivo maturation from selection of a pre-existing engrafting subset; stem-enriched subset profiling, single-cell methylome approaches, and clonal tracking would refine this resolution. Integrating these approaches would determine whether young-like methylomes correlate with preservation of clonal diversity and reduced myeloid bias, a key determinant of functional aging in hematopoiesis [Pang et al, 2011; Ross et al, 2024].

In summary, we show that polyclonal, donor-diverse iPSC-derived HSC grafts can achieve a primary-HSC-like epigenetic state after long-term engraftment while preserving young-like epigenetic features, even through serial transplantation. These results support the concept that definitive HSC identity and epigenetic aging are dissociable, opening a path to engineer transplantable hematopoietic grafts that combine adult function with youthful molecular reserve.

## Materials and methods

### iPSC culture and maintenance

Human iPSC lines were purchased from Cedars-Sinai Biomanufacturing Center or Pluristyx (Table 1). All lines were derived from healthy adult donors and were maintained in culture on flasks coated for 1 hour with 2.4 µL/mL iMatrix-511 (Takara Bio #892011) in PBS (Cytiva #SH30256.01). Cells were kept in mTeSR Plus media (STEMCELL #100-1130) which was supplemented with 10 uM ROCK inhibitor Thiazovivin (Tocris #3845) for 24 hours after passaging.

**Table 1.**
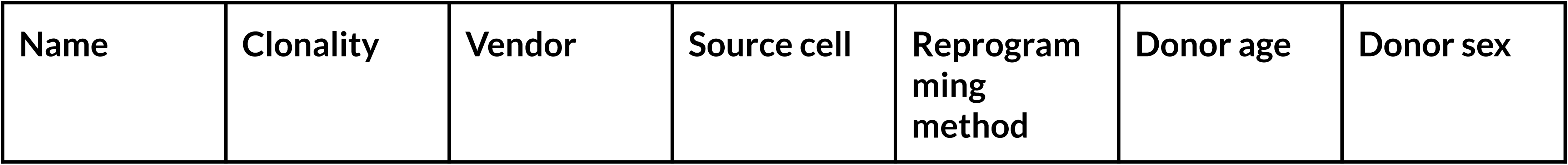
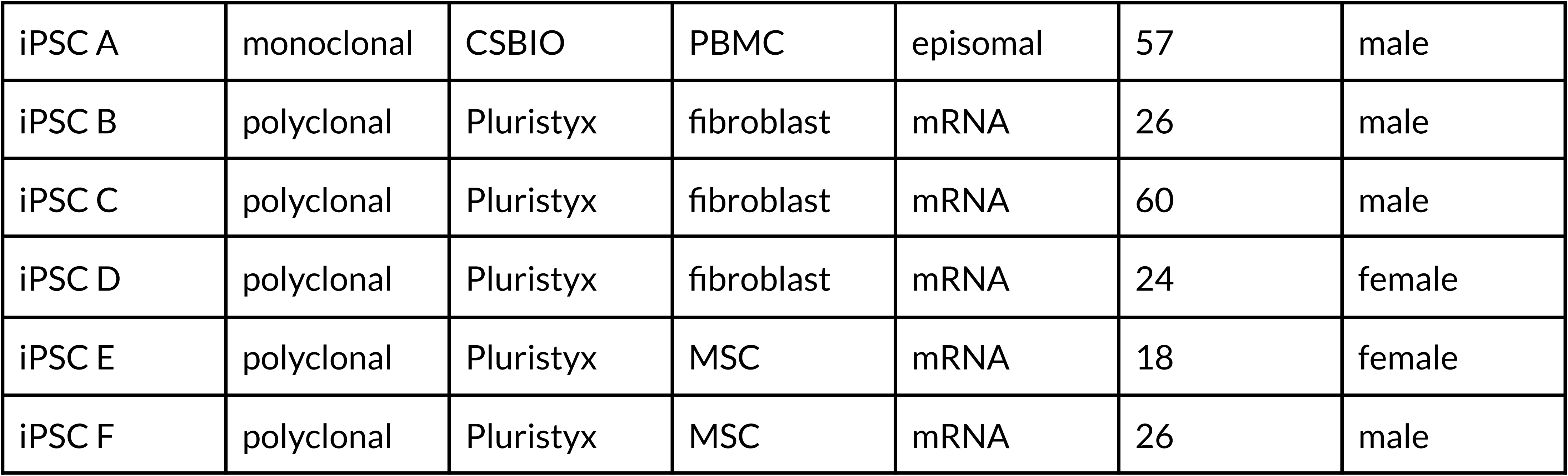
Human iPSC lines.

| Name | Clonality | Vendor | Source cell | Reprogram<br>ming<br>method | Donor age | Donor sex |
| --- | --- | --- | --- | --- | --- | --- |
| iPSC A | monoclonal | CSBIO | PBMC | episomal | 57 | male |
| iPSC B | polyclonal | Pluristyx | fibroblast | mRNA | 26 | male |
| iPSC C | polyclonal | Pluristyx | fibroblast | mRNA | 60 | male |
| iPSC D | polyclonal | Pluristyx | fibroblast | mRNA | 24 | female |
| iPSC E | polyclonal | Pluristyx | MSC | mRNA | 18 | female |
| iPSC F | polyclonal | Pluristyx | MSC | mRNA | 26 | male |

### iHSC differentiations

Hematopoietic differentiation was performed using the swirling embryoid body method [Ng et al, 2024]. Human iPSC cultures were dissociated using 0.5 mM EDTA (Invitrogen #15575-038) solution for 3-5 min and resuspended in differentiation base media (Table 2) for cell counts. 1 × 10^6^ dissociated cells were transferred to each non-treated 60-mm dish (Greiner #628161) in 5 mL of differentiation base media supplemented with 4 uM CHIR99021 (Tocris #4423), 30 ng/mL Activin A (R&D Systems #AFL338), 20 ng/mL bFGF (R&D Systems #FGFB), and 1 uM Thiazovivin (Tocris #3845). Dishes were placed on a Benchmark BT4001 orbital shaker rotating at 60 rpm in a 5% CO_2_ incubator at 37°C. Differentiation media was replaced with 6 mL base media with additional cytokines on day 1 and subsequently every other day (days 3, 5, 7, 9, 11, and 13) during the 15-day differentiation with growth factors as previously described [Ng et al, 2024] (Table 3).

**Table 2.** Components of iPSC→iHSC differentiation base media.

| Component | Vendor | Cat # | Stock concentration | Final concentration |
| --- | --- | --- | --- | --- |
| IMDM | Gibco | 21056023 | 1x | 0.5x |
| F12 | Gibco | 11765047 | 1x | 0.5x |
| 10% PVA in dH<sub>2</sub>O | Sigma Aldrich | P8136-250G | 10% | 0.06% |
| Glutamax | Life Technologies | 35050061 | 100x | 1x |
| L-AA | Tocris | A4544-100G | 10 mg/mL | 50 μg/mL |
| AA2P | Tocris | A8960-5G | 10 mg/mL | 50 μg/mL |
| ITSE/ITX | Thermo Fisher | 51500056 | 100x | 1x |
| NEAA | Life Technologies | 11140050 | 100x | 1x |
| Synthechol | Sigma Aldrich | S5442-2ML | 500x | 0.15x |

**Table 3.** Cytokines and small molecules for iPSC→iHSC differentiation.

| Component | Vendor | Cat # | Final concentration |
| --- | --- | --- | --- |
| BMP4 | R&D Systems | AFL314E | 20 ng/mL (days 3-5)<br>2 ng/mL (days 5-11) |
| CHIR99021 | Tocris | 4423 | 4 uM (day 0-1)<br>3 uM (days 1-3) |
| Activin A | R&D Systems | AFL338 | 30 ng/mL (day 0-1) |
| bFGF | R&D Systems | BT-FGFB-AFL | 20 ng/mL (days 0-15) |
| Thiazovivin | Tocris | 3845 | 1 uM (day 0-1) |
| SB431542 | Tocris | 1614 | 4 uM (days 1-3) |
| VEGF | R&D Systems | BT-VEGF-AFL | 50 ng/mL (days 1-3)<br>150 ng/mL (days 3-7) |
| RETA | Sigma Aldrich | R0635 | 100 nM (days 1-3, 5-15)<br>2 uM (days 3-5) |
| SCF | R&D Systems | BT-SCF-AFL | 10 ng/mL (days 3-15) |
| TPO | R&D Systems | 288E-GMP | 10 ng/mL (days 11-15) |
| UM729 | STEMCELL | 72332 | 250 nM (days 11-15) |
| IGF1 | R&D Systems | 291-GMP | 10 ng/mL (days 3-11)<br>5 ng/mL (days 11-15) |
| IGF2 | R&D Systems | 292-G2 | 10 ng/mL (days 3-11)<br>5 ng/mL (days 11-15) |
| SR1 | Tocris | 7086 | 100 nM (days 11-15) |

Blood cells began to be shed into suspension by the embryoid bodies after 9 days of differentiation. On day 15, cultures were harvested and suspension cells were separated from the swirling EBs by passing the cell suspension through 40 um filters (Corning #431750). The day 15 suspension cells (iHSCs) were analyzed by flow cytometry and were cryopreserved in 10% DMSO media, CryoStor CS10 (STEMCELL #07930), for downstream assays.

### Quality control

Karyotyping of iPSC lines was performed by the respective vendors, Cedars-Sinai Biomanufacturing Center or Pluristyx (Table 1). Day 15 iHSCs were processed by IDEXX Bioanalytics for their iCS-digital assay for copy number variant (CNV) genomic testing, as well as for short tandem repeat (STR) analysis and mycoplasma testing. iHSCs were confirmed to pass all quality control assays before downstream assays.

### Flow cytometry

At days 3, 5, 7, 9, 11, 13 and 15 of iPSC to iHSC differentiation, dissociated EBs and suspension hematopoietic cells were analyzed by flow cytometry using a CytoFLEX LX Flow Cytometer (Beckman Coulter). Flow cytometry was also carried out on suspension cells from days 8, 9, 10, and 11 of platelet and neutrophil differentiation from HSCs. EBs were dissociated by 1000 rpm incubation with 2 mg/mL collagenase type II (Thermo Scientific #17101-015) at 42°C for 30 minutes followed by mechanical dissociation by passing through a 25-gauge needle and resuspension in cold PBS (Cytiva #SH30256.01) supplemented with 2% fetal bovine serum (Corning #35-016-CV) and 2 mM EDTA (Invitrogen #15575-038), referred to henceforth as FACS buffer. Suspension cells were analyzed separately from dissociated EB cells.

Transplanted mouse tissues were analyzed for human hematopoietic cell engraftment with flow cytometry using a CytoFLEX LX Flow Cytometer. Red blood cell lysis of peripheral blood samples was performed by incubating 200 uL of blood with 10 mL of ammonium-chloride-potassium (ACK) lysing buffer (Quality Biological #118-156-101) for 15 min at room temperature followed by wash and resuspension in FACS buffer. Hematopoietic cells were collected from the bone marrow (of the mouse tibia, femur, and ilium) and spleen by mechanical dissociation and filtering. Bone marrow single cell samples were sometimes split for multiple antibody panels.

Fluorescent-conjugated antibodies were used to identify cell populations (Table 4). All samples were incubated with the appropriate dilution of antibodies in a volume of 100 uL of FACS buffer for 15 min at 4°C, washed with FACS buffer, and resuspended in 200 uL of FACS buffer for flow cytometric analysis. Gates were defined using fluorescence minus one stainings.

**Table 4.** Antibodies for flow cytometry.

| Antigen | Fluorophore | Dilution | Vendor | Cat # |
| --- | --- | --- | --- | --- |
| Mouse CD45 | Pacific Blue | 1:100 | BioLegend | 103126 |
| Human CD45 | FITC | 1:100 | BioLegend | 982316 |
| Human CD43 | PE | 1:100 | BioLegend | 343204 |
| Human CD19 | Brilliant Violet 605 | 1:100 | BioLegend | 302244 |
| Human CD33 | Brilliant Violet 785 | 1:100 | BioLegend | 303428 |
| Human CD3 | Alexa Fluor 700 | 1:100 | BioLegend | 344822 |
| Human CD235a | APC | 1:200 | BioLegend | 551336 |
| Human CD133 | PE | 1:100 | BioLegend | 393904 |
| Human CD34 | PE-Cyanine7<br>PE | 1:100 | BioLegend | 343516 |
| Human CD38 | Brilliant Violet<br>421 | 1:100 | BioLegend | 356618 |
| Human Lineage | Brilliant Violet<br>510 | 1:100 | BioLegend | 348807 |
| Human CD41 | APC-Cyanine7 | 1:100 | BioLegend | 303716 |
| Human CD42b | Brilliant Violet<br>650 | 1:100 | BioLegend | 303926 |
| Human CD14 | PE/Dazzle 594 | 1:100 | BioLegend | 301852 |
| Human CD15 | PE-Cyanine7 | 1:100 | BioLegend | 323030 |
| Human CD16 | Brilliant Violet<br>605 | 1:100 | BioLegend | 302040 |
| Human CD66b | Alexa Fluor 700 | 1:100 | BioLegend | 305114 |
| Human CXCR4 | APC | 1:100 | BioLegend | 306510 |
| Human KDR | PE-Cyanine7 | 1:100 | BioLegend | 393008 |
| Human CD73 | Brilliant Violet<br>421 | 1:100 | BioLegend | 344008 |
| Human CD90 | APC | 1:100 | BioLegend | 328114 |
| Human CD201 | PE | 1:100 | BioLegend | 351904 |
| Human PAC-1 | FITC | 1:100 | BD | 340507 |
| Human CD62p | APC | 1:100 | BD | 550888 |
| 7-AAD | n/a | 1:100 | BioLegend | 420404 |
| DAPI | n/a | 1:1000 | Miltenyi Biotec | 130-111-570 |

### Primary HSC samples

Mixed donor human umbilical cord blood (CB) CD34+ cells were purchased from STEMCELL Technologies (#70008) and thawed, washed, and resuspended for transplantation as described below (see Transplantation experiments section).

Adult primary HSC controls were isolated from mobilized peripheral blood (mPB) leukopaks or CD34+ vials purchased from Charles River (Table 5). For leukopak donors, CD34+ cells were isolated from leukopaks using anti-human CD34 antibody-conjugated magnetic beads (Miltenyi Biotec #130-046-702) according to the manufacturer’s protocol and cryopreserved for downstream assays.

**Table 5.**
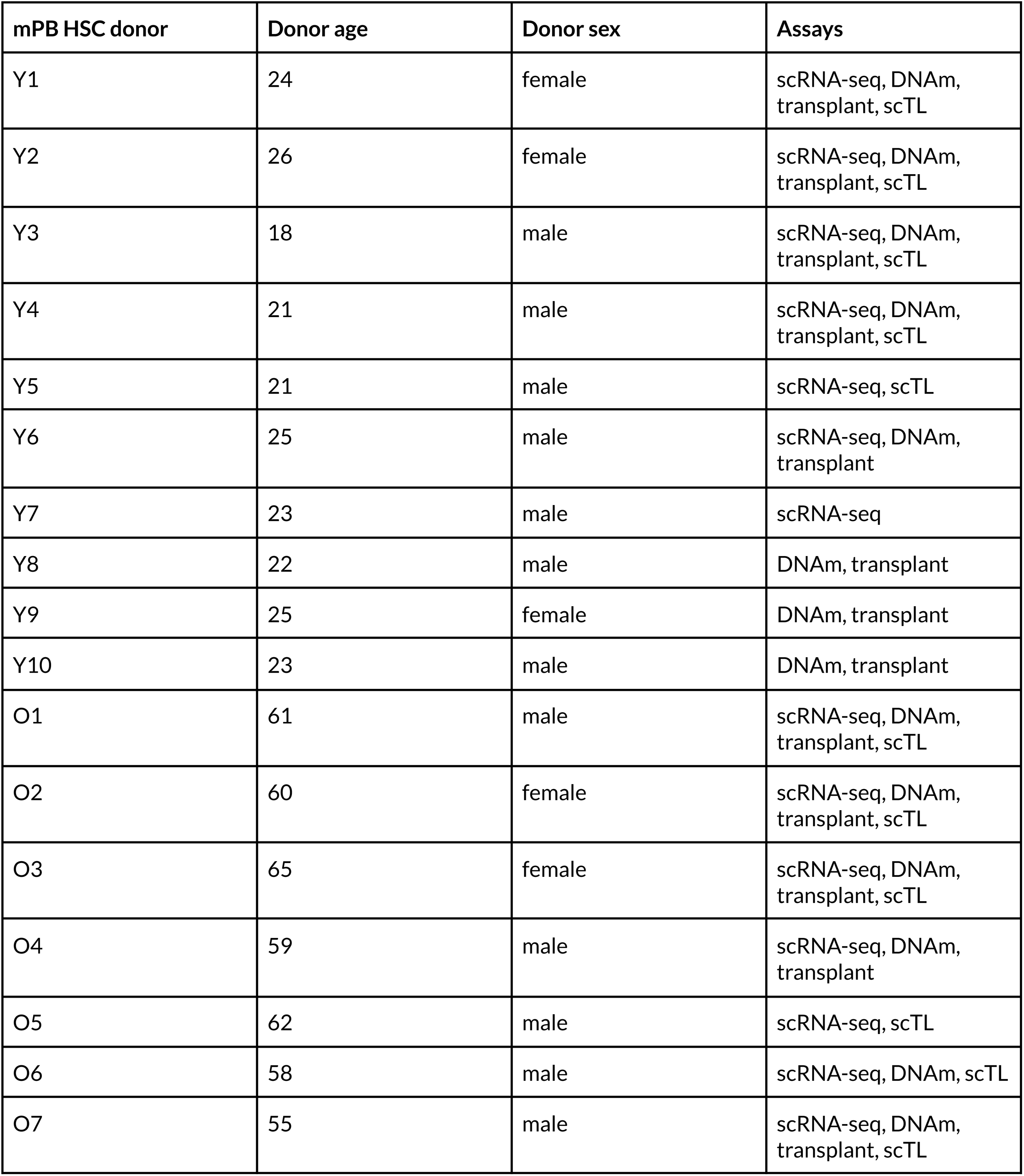
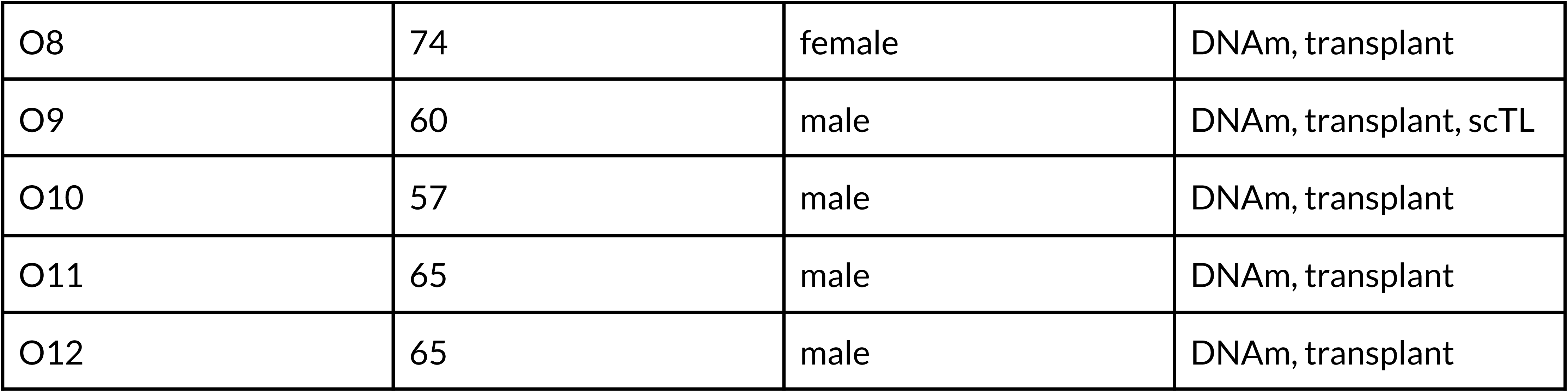
Adult mobilized peripheral blood donors.

### Animals

NOD.Cg-*Kit^W-41J^ Tyr^+^ Prkdc^scid^ Il2rg^tm1Wjl^*/ThomJ (NBSGW) mice [McIntosh et al, 2015] were sourced from The Jackson Laboratory, and only female mice were used. Mice were housed up to five animals per cage, and kept on a 12 hour light cycle. All experiments using mice were performed in accordance with the Retro Biosciences guidelines for the use of laboratory animals and were approved by the Institutional Animal Care and Use Committees.

### Transplantation experiments

Cryopreserved iHSCs or primary HSCs (CB or mPB CD34+ cells) were thawed, washed, and resuspended in differentiation base media (Table 2) + 2% FBS for primary transplantations. All primary HSCs were CD34+ enriched and iHSCs were transplanted if > 70% CD34+CD45+. Female NBSGW mice aged between 7 and 13 weeks were injected retro-orbitally, each with 5 × 10^6^ iHSCs, 5 × 10^4^ CB HSCs, or 5 × 10^5^ – 1 × 10^6^ mPB HSCs. Peripheral blood samples (up to 200 uL) were collected from transplanted mice every 4 weeks via saphenous bleeds. At 20 weeks post-transplant, hind legs, spleens, and peripheral blood were collected for terminal engraftment analysis. Tissues were processed into single-cell suspensions and cells were analyzed via flow cytometry (see Flow cytometry section) for human cell surface antigens indicative of erythroid, myeloid, B, T, and stem cell compartments (Table 4). Residual bone marrow samples from engrafted mice were cryopreserved for further assays. Bone marrow samples from selected primary transplant groups were collected and transplanted on the same day, without cryopreservation, into young female NBSGW mice via retro-orbital injection (up to 10 × 10^6^ whole bone marrow cells per mouse). Secondary transplant groups were monitored in the same manner as in the primary transplants.

### Transcriptional profiling with scRNA-seq

Single-cell RNA-sequencing was performed on 13 lots of independently differentiated iHSCs from 6 iPSC lines (Table 1) alongside primary mPB and CB HSCs and 2 iPSC samples. Cryopreserved cell samples were thawed and libraries were prepared with the 10x Genomics fixed scRNA-seq kit according to the manufacturer’s protocol and sequenced on Illumina platforms.

Raw sequencing data were processed using Cell Ranger with alignment to the GRCh38 human reference genome to generate gene-by-cell count matrices. Ambient RNA contamination was removed using CellBender. Subsequent analyses were performed in Python using Scanpy and scvi-tools. Cells were filtered based on gene count, UMI count, mitochondrial transcript fraction, and doublet detection criteria. Batch correction and integration across donors were performed using scVI and UMAP embeddings and clustering were computed from the scVI latent space. Cell type annotation was performed based on canonical marker gene expression and comparison to published human HSC atlases [Calvanese et al, 2022; Ng et al, 2024].

### DNA methylation profiling

Genomic DNA (gDNA) was extracted from cell samples with the DNeasy Blood and Tissue Kit (QIAGEN #69506). For samples at early stages of differentiation, EBs were dissociated as described above (see Flow cytometry section) and gDNA was extracted from the resulting single cell suspension. For engrafted tissue samples, cryopreserved bone marrow was thawed and anti-human CD45 antibody-conjugated magnetic beads (Miltenyi Biotec #130-045-801) were used to isolate human hematopoietic cells for gDNA extraction.

gDNA samples were then further processed at TruDiagnostic (Lexington, KY) and run on Infinium MethylationEPIC v2.0 BeadChip arrays (Illumina) for methylation profiling at 929,386 CpG sites [Kaur et al, 2023]. Raw IDAT files were processed by TruDiagnostic using a single-sample Noob (ssNoob) normalization pipeline implemented in the minfi R package [Aryee et al, 2014; Fortin et al, 2014]. Briefly, preprocessing included background subtraction using out-of-band probes, dye-bias correction, zero-intensity imputation to prevent division-by-zero artifacts, and conversion of corrected methylated and unmethylated intensities into beta values with denominator offset. Quality control metrics included signal intensity thresholds and exclusion of poorly performing probes as described previously [Kaur et al, 2023; Lin et al, 2026].

DNA methylation age was calculated by TruDiagnostic from the normalized beta values using the PC-based versions [Higgins-Chen et al, 2022] of the Horvath multi-tissue clock [Horvath, 2013], the Hannum clock [Hannum et al, 2013], and the PhenoAge clock [Levine et al, 2018]. Shannon entropy of age-associated sites was calculated as described in Hannum et al, 2013. Immune cell type deconvolution was performed with the Luo et al, 2023 reference matrix. DunedinPACE [Belsky et al, 2022] and SystemsAge [Sehgal et al, 2025] scores were also provided.

Normalized beta values and epigenetic clock outputs and scores were returned to our laboratory for downstream analysis. Gene promoter methylation values were calculated as the mean normalized beta value across CpG sites annotated to promoter regions (TSS200 and TSS1500). Differential methylation analysis at the promoter level was performed using a two-sided t-test implemented in Scanpy comparing specified cell-type groups. P values were adjusted for multiple testing, and promoters were considered differentially methylated if they met a false discovery rate (FDR) < 0.05, absolute log fold change (|LFC|) > 1, and |z-score| > 2.

### Telomere length measurements

Single-cell telomere length measurements were performed by Life Length Laboratories (Madrid, Spain) using their patented high-throughput quantitative fluorescence in situ hybridization–based Telomere Analysis Technology (TAT), as previously described [Canela et al, 2007; De Pedro et al, 2020]. Cryopreserved cell samples were shipped on dry ice and processed according to Life Length standard operating procedures [De Pedro et al, 2020]. Image acquisition and analysis was performed on a High Content Screening Opera Phenix System (Revvity) using the SIMA software, Version 1.2 (Revvity). For each sample, five technical replicates (independently seeded and imaged wells) were processed, and telomere length distributions were calculated from >10,000 individual telomeric measurements. Median telomere length values and critically short telomere fractions of iHSCs and primary HSCs were compared with a two-sided Wilcoxon rank-sum test.

### Lineage differentiations and functional assays

iHSCs and young and old mPB HSCs were thawed into non-tissue-culture-treated 6-well plates at a density of 5 × 10^5^ cells/well. For differentiations into neutrophil progenitors, HSCs were seeded with differentiation base media with 100 ng/mL G-CSF (STEMCELL #78012), 50 ng/mL SCF (R&D Systems #BT-SCF-AFL), and 50 ng/mL IL-3 (R&D Systems #203-IL). Cells were expanded to more wells as necessary and IL-3 was removed after day 3. For differentiations into platelet progenitors, HSCs were seeded with 100 ng/mL TPO (R&D Systems #288E-GMP) and 50 ng/mL SCF and similarly expanded as necessary. Both platelets and neutrophils were harvested and cryopreserved on day 11.

| Component | Vendor | Cat # | Final concentration |
| --- | --- | --- | --- |
| G-CSF | STEMCELL | 78012 | 100 ng/mL |
| SCF | R&D Systems | BT-SCF-AFL | 50 ng/mL |
| IL-3 | R&D Systems | 203-IL | 50 ng/mL |
| TPO | R&D Systems | 288E-GMP | 100 ng/mL |

Neutrophils were thawed and recovered overnight in base media with 100 ng/mL G-CSF and 50 ng/mL SCF. Neutrophils were then seeded at 5 × 10^4^ cells/well in 96-well plates, treated with pHrodo Red *E. coli* BioParticles (Thermo Fisher #A10025), and incubated in an IncuCyte (Sartorius) for imaging.

HSC-derived platelets were thawed and recovered overnight in differentiation base media with 100 ng/mL TPO and 50 ng/mL SCF. Platelets were then seeded at in 96-well plates and treated with 5 uM ADP (Sigma-Aldrich #A2754) or 10 uM TRAP-6 (Tocris #3497) for 20 min at room temperature and analyzed for PAC-1 or CD62P expression via flow cytometry.

### Cell counts

Routine cell counts and viability measurements were performed with the NucleoCounter NC-200 (ChemoMetec).

### Cryopreservation

Cells were cryopreserved in 10% DMSO media, CryoStor CS10 (STEMCELL #07930), using Mr. Frosty Freezing Containers (Thermo Scientific #5100-0001) at -80°C. Vials were then transferred for long-term storage in liquid nitrogen tanks.

---

**FIGURE S1.**
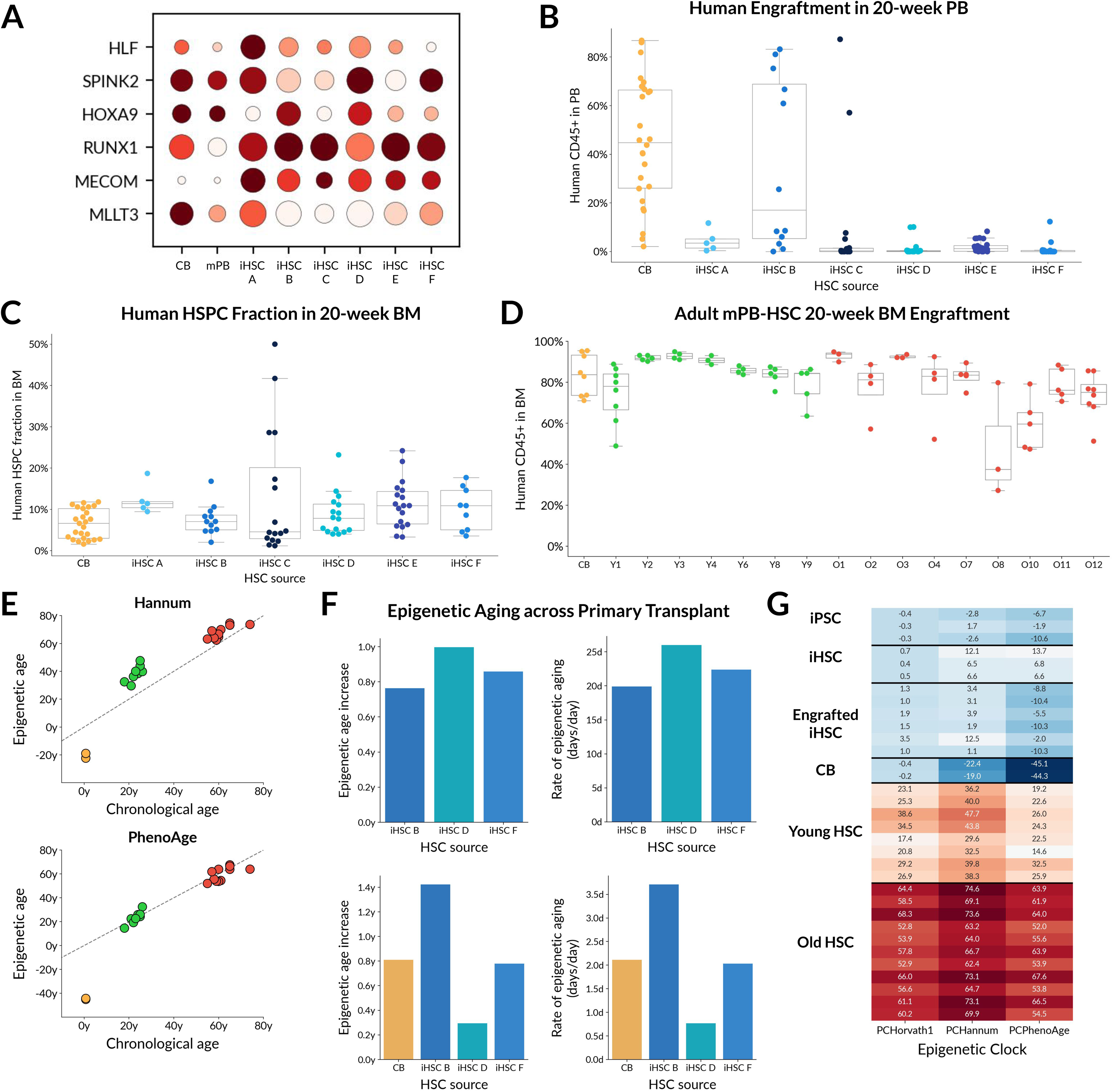
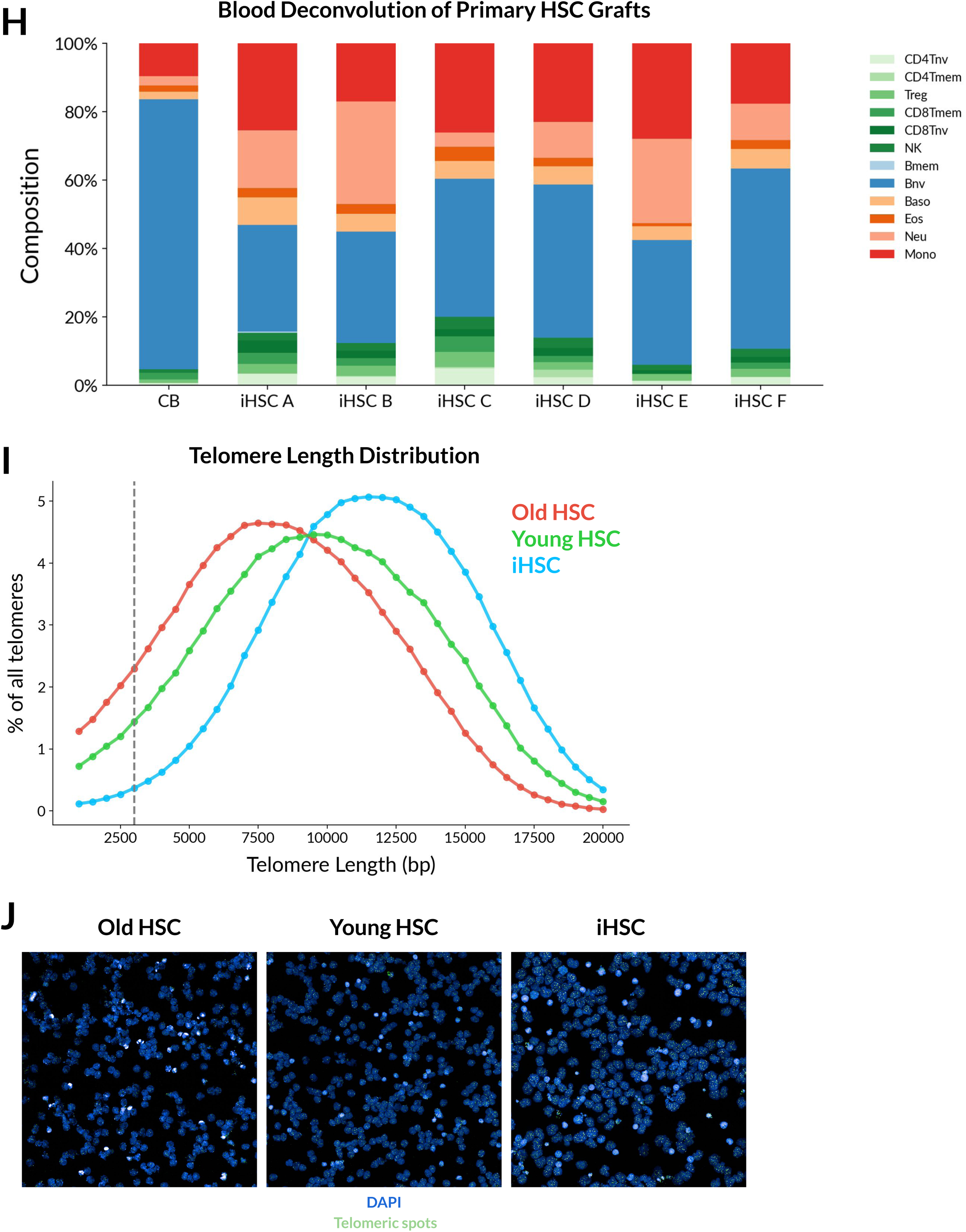
**(A)** Dot plot of HSC signature transcription factors in cord blood (CB), mobilized peripheral blood (mPB), and iHSCs from six donors measured by single-cell RNA-sequencing. Dot size indicates the fraction of cells expressing each gene and color indicates relative expression level. **(B)** Human CD45+ chimerism in peripheral blood of NBSGW recipients at 20 weeks after transplantation with CB CD34+ cells or iHSCs from the indicated donor lines. **(C)** Frequency of phenotypic human HSPCs (CD45+CD34+CD38–) in 20-week bone marrow grafts after transplantation with CB CD34+ cells or iHSCs. **(D)** Human CD45+ bone marrow chimerism in NBSGW recipients transplanted with adult mPB CD34+ HSC controls from young and old donors. **(E)** Hannum and PhenoAge epigenetic clock predictions for primary fetal/CB, young adult, and old adult CD34+ HSC controls plotted against chronological age. **(F)** Epigenetic age increase and rate of epigenetic aging across primary transplantation for CB and longitudinally profiled iHSC grafts. **(G)** Heatmap of PC-based Horvath, Hannum, and PhenoAge clock predictions across iPSCs, day 15 iHSCs, engrafted iHSC-derived cells, CB, young primary HSCs, and old primary HSCs. **(H)** DNA methylation-based blood cell deconvolution of primary CB and iHSC-derived grafts after long-term engraftment. **(I)** Telomere length distribution of day 15 iHSCs compared with young and old adult primary HSCs. The dashed line indicates the 3-kb threshold used to define critically short telomeres. **(J)** Representative HT Q-FISH images of old primary HSCs, young primary HSCs, and iHSCs. DAPI marks nuclei and telomeric probe signal marks telomere foci.

**FIGURE S2.**
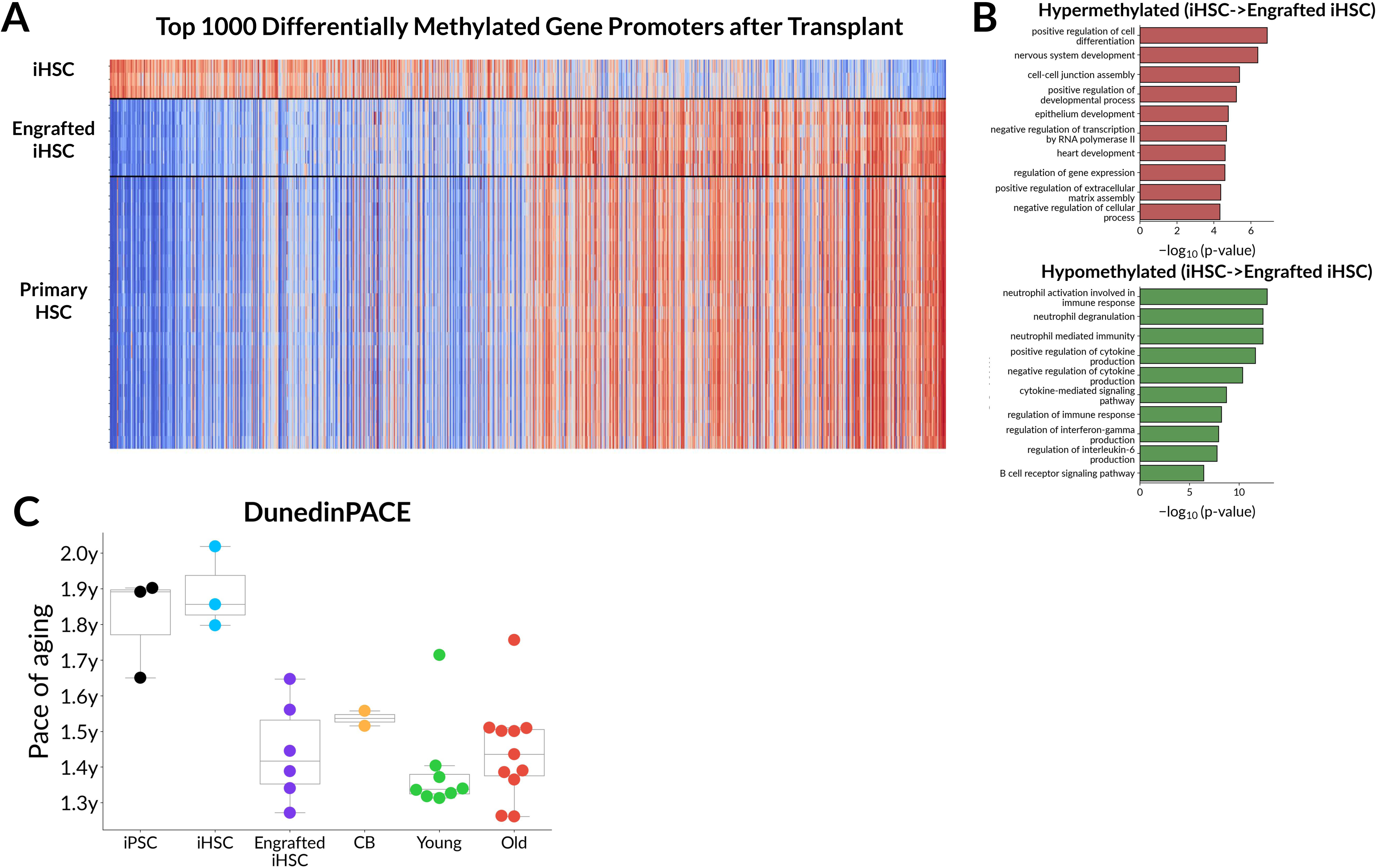
**(A)** Heatmap of the top 1,000 differentially methylated gene promoters between pre-transplant iHSCs and 20-week engrafted iHSC-derived human cells, shown alongside primary HSC controls. Engrafted iHSC-derived cells shift toward the promoter methylation state of primary HSCs. **(B)** Gene ontology enrichment for promoters that became hypermethylated or hypomethylated in engrafted iHSC-derived cells relative to pre-transplant iHSCs. Hypermethylated promoters were enriched for developmental and regulatory processes, while hypomethylated promoters were enriched for immune, cytokine, and neutrophil-associated programs. **(C)** DunedinPACE scores across iPSCs, day 15 iHSCs, engrafted iHSC-derived cells, CB, young primary HSCs, and old primary HSCs. DunedinPACE was elevated in iPSCs and pre-transplant iHSCs and decreased after long-term engraftment toward primary hematopoietic cell values.

**FIGURE S3.**
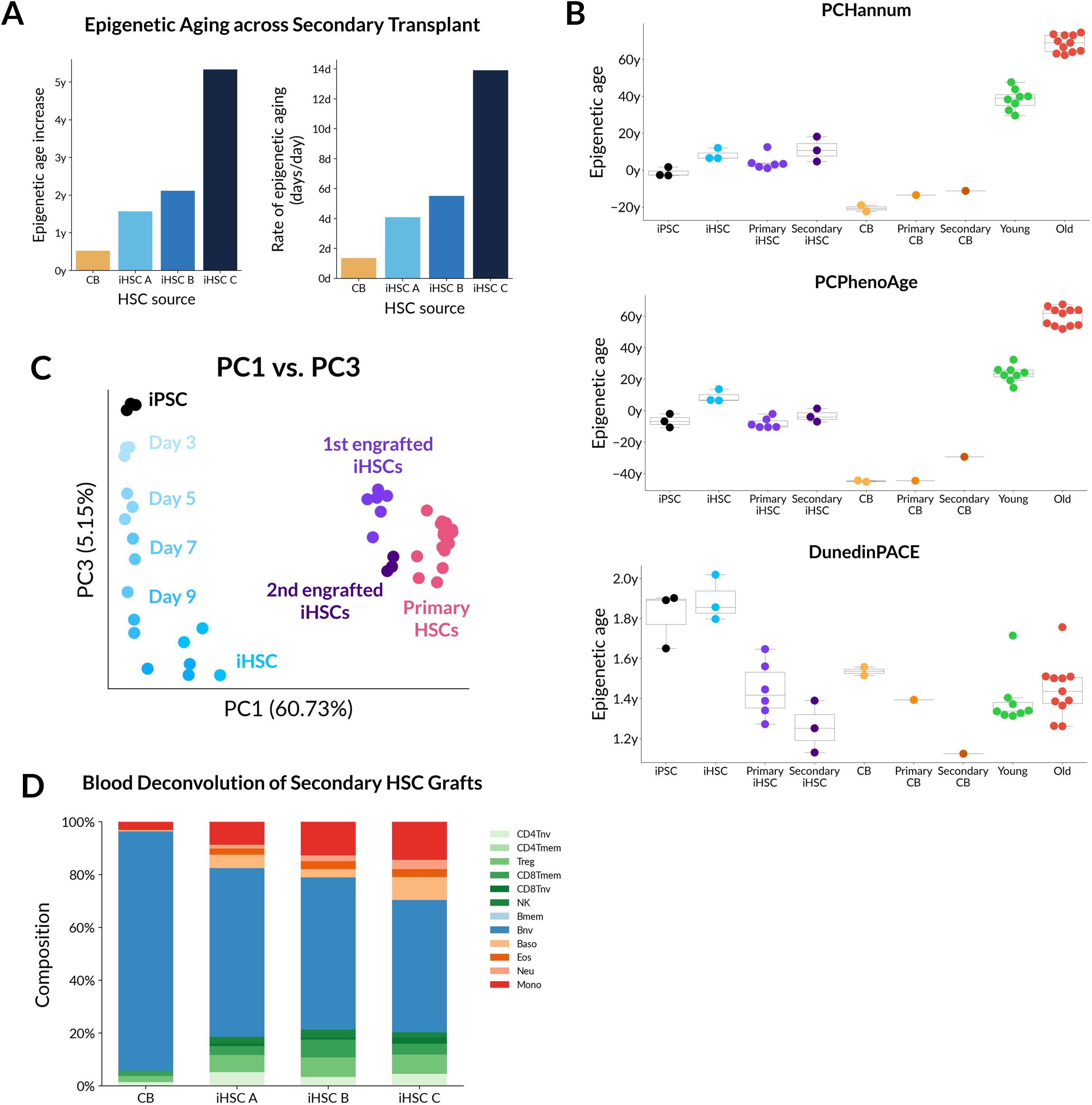
**(A)** Epigenetic age increase and rate of epigenetic aging across secondary transplantation for CB and iHSC-derived grafts, calculated from Horvath clock predictions. **(B)** PC-based Hannum, PC-based PhenoAge, and DunedinPACE estimates across iPSCs, pre-transplant iHSCs, primary engrafted iHSC-derived cells, secondary engrafted iHSC-derived cells, primary and secondary CB grafts, young primary HSCs, and old primary HSCs. Secondary engrafted iHSC-derived cells retain young epigenetic age estimates without reversion to donor age. **(C)** Principal component analysis of genome-wide DNA methylation profiles across iPSC-to-iHSC differentiation, primary engraftment, secondary engraftment, and primary HSC controls. Serially engrafted iHSC-derived cells remain close to primary engrafted iHSC-derived cells and primary HSCs in methylation space. **(D)** DNA methylation-based blood cell deconvolution of secondary CB and iHSC-derived grafts.

## References

1. Abid, M. B., Estrada-Merly, N., Zhang, M. J., Chen, K., Allan, D., Bredeson, C., Sabloff, M., Guru Murthy, G. S., Badar, T., Hashmi, S., Aljurf, M., Litzow, M. R., Kebriaei, P., Hourigan, C. S., & Saber, W. (2023). Impact of Donor Age on Allogeneic Hematopoietic Cell Transplantation Outcomes in Older Adults with Acute Myeloid Leukemia. Transplantation and Cellular Therapy, 29(9), 578.e1–578.e9. 10.1016/J.JTCT.2023.06.020

2. Aryee, M. J., Jaffe, A. E., Corrada-Bravo, H., Ladd-Acosta, C., Feinberg, A. P., Hansen, K. D., & Irizarry, R. A. (2014). Minfi: a flexible and comprehensive Bioconductor package for the analysis of Infinium DNA methylation microarrays. Bioinformatics, 30(10), 1363–1369. 10.1093/BIOINFORMATICS/BTU049

3. Aubert, G., Baerlocher, G. M., Vulto, I., Poon, S. S., & Lansdorp, P. M. (2012). Collapse of Telomere Homeostasis in Hematopoietic Cells Caused by Heterozygous Mutations in Telomerase Genes. PLOS Genetics, 8(5), e1002696. 10.1371/JOURNAL.PGEN.1002696

4. Beerman, I., Bock, C., Garrison, B. S., Smith, Z. D., Gu, H., Meissner, A., & Rossi, D. J. (2013). Proliferation-Dependent Alterations of the DNA Methylation Landscape Underlie Hematopoietic Stem Cell Aging. Cell Stem Cell, 12(4), 413–425. 10.1016/J.STEM.2013.01.017

5. Belsky, D. W., Caspi, A., Corcoran, D. L., Sugden, K., Poulton, R., Arseneault, L., Baccarelli, A., Chamarti, K., Gao, X., Hannon, E., Harrington, H. L., Houts, R., Kothari, M., Kwon, D., Mill, J., Schwartz, J., Vokonas, P., Wang, C., Williams, B., & Moffitt, T. E. (2022). DunedinPACE, A DNA methylation biomarker of the Pace of Aging. eLife, 11. 10.7554/ELIFE.73420

6. Bocklandt, S., Lin, W., Sehl, M. E., Sánchez, F. J., Sinsheimer, J. S., Horvath, S., & Vilain, E. (2011). Epigenetic Predictor of Age. PLOS ONE, 6(6), e14821. 10.1371/JOURNAL.PONE.0014821

7. Calvanese, V., Capellera-Garcia, S., Ma, F., Fares, I., Liebscher, S., Ng, E. S., Ekstrand, S., Aguadé-Gorgorió, J., Vavilina, A., Lefaudeux, D., Nadel, B., Li, J. Y., Wang, Y., Lee, L. K., Ardehali, R., Iruela-Arispe, M. L., Pellegrini, M., Stanley, E. G., Elefanty, A. G., … Mikkola, H. K. A. (2022). Mapping human haematopoietic stem cells from haemogenic endothelium to birth. Nature 2022 604:7906, 604(7906), 534–540. 10.1038/s41586-022-04571-x

8. Canela, A., Vera, E., Klatt, P., & Blasco, M. A. (2007). High-throughput telomere length quantification by FISH and its application to human population studies. Proceedings of the National Academy of Sciences of the United States of America, 104(13), 5300–5305. 10.1073/PNAS.0609367104;PAGE:STRING:ARTICLE/CHAPTER

9. Cimato, T. R., Conway, A., Nichols, J., & Wallace, P. K. (2019). CD133 expression in circulating hematopoietic progenitor cells. Cytometry Part B - Clinical Cytometry, 96(1), 39–45. 10.1002/CYTO.B.21732;PAGE:STRING:ARTICLE/CHAPTER

10. Copelan, E. A. (2006). Hematopoietic Stem-Cell Transplantation. New England Journal of Medicine, 354(17), 1813–1826. 10.1056/NEJMRA052638

11. de Pedro, N., Díez, M., García, I., García, J., Otero, L., Fernández, L., García, B., González, R., Rincón, S., Pérez, D., Rodríguez, E., Segovia, E., & Najarro, P. (2020). Analytical Validation of Telomere Analysis Technology® for the High-Throughput Analysis of Multiple Telomere-Associated Variables. Biological Procedures Online 2020 22:1, 22(1), 2-. 10.1186/S12575-019-0115-Z

12. Dezern, A. E., Franklin, C., Tsai, H. L., Imus, P. H., Cooke, K. R., Varadhan, R., & Jones, R. J. (2021). Relationship of donor age and relationship to outcomes of haploidentical transplantation with posttransplant cyclophosphamide. Blood Advances, 5(5), 1360–1368. 10.1182/BLOODADVANCES.2020003922

13. Ditadi, A., Sturgeon, C. M., Tober, J., Awong, G., Kennedy, M., Yzaguirre, A. D., Azzola, L., Ng, E. S., Stanley, E. G., French, D. L., Cheng, X., Gadue, P., Speck, N. A., Elefanty, A. G., & Keller, G. (2015). Human definitive haemogenic endothelium and arterial vascular endothelium represent distinct lineages. Nature Cell Biology, 17(5), 580–591. 10.1038/NCB3161;TECHMETA

14. Flach, J., Bakker, S. T., Mohrin, M., Conroy, P. C., Pietras, E. M., Reynaud, D., Alvarez, S., Diolaiti, M. E., Ugarte, F., Forsberg, E. C., le Beau, M. M., Stohr, B. A., Méndez, J., Morrison, C. G., & Passegué, E. (2014). Replication stress is a potent driver of functional decline in ageing haematopoietic stem cells. Nature 2014 512:7513, 512(7513), 198–202. 10.1038/nature13619

15. Fortin, J. P., Triche, T. J., & Hansen, K. D. (2017). Preprocessing, normalization and integration of the Illumina HumanMethylationEPIC array with minfi. Bioinformatics, 33(4), 558–560. 10.1093/BIOINFORMATICS/BTW691

16. Frobel, J., Hemeda, H., Lenz, M., Abagnale, G., Joussen, S., Denecke, B., Šarić, T., Zenke, M., & Wagner, W. (2014). Epigenetic Rejuvenation of Mesenchymal Stromal Cells Derived from Induced Pluripotent Stem Cells. Stem Cell Reports, 3(3), 414–422. 10.1016/J.STEMCR.2014.07.003

17. Gadalla, S. M., Aubert, G., Wang, T., Haagenson, M., Spellman, S. R., Wang, L., Katki, H. A., Savage, S. A., & Lee, S. J. (2018). Donor telomere length and causes of death after unrelated hematopoietic cell transplantation in patients with marrow failure. Blood, 131(21), 2393–2398. 10.1182/BLOOD-2017-10-812735

18. Gadalla, S. M., Katki, H. A., Lai, T. P., Auer, P. L., Dagnall, C. L., Bupp, C., Hutchinson, A. A., Anderson, J. J., Mendez, K. J. W., Spellman, S. R., Stewart, V., Savage, S. A., Lee, S. J., Levine, J. E., Saber, W., & Aviv, A. (2025). Donor telomeres and their magnitude of shortening post-allogeneic haematopoietic cell transplant impact survival for patients with early-stage leukaemia or myelodysplastic syndrome. eBioMedicine, 114, 105641. 10.1016/j.ebiom.2025.105641

19. Gadalla, S. M., Wang, T., Haagenson, M., Spellman, S. R., Lee, S. J., Williams, K. M., Wong, J. Y., de Vivo, I., & Savage, S. A. (2015). Association Between Donor Leukocyte Telomere Length and Survival After Unrelated Allogeneic Hematopoietic Cell Transplantation for Severe Aplastic Anemia. JAMA, 313(6), 594–602. 10.1001/JAMA.2015.7

20. Garagnani, P., Bacalini, M. G., Pirazzini, C., Gori, D., Giuliani, C., Mari, D., di Blasio, A. M., Gentilini, D., Vitale, G., Collino, S., Rezzi, S., Castellani, G., Capri, M., Salvioli, S., & Franceschi, C. (2012). Methylation of ELOVL2 gene as a new epigenetic marker of age. Aging Cell, 11(6), 1132–1134. 10.1111/ACEL.12005;WGROUP:STRING:PUBLICATION

21. Görgens, A., Radtke, S., Möllmann, M., Cross, M., Dürig, J., Horn, P. A., & Giebel, B. (2013). Revision of the Human Hematopoietic Tree: Granulocyte Subtypes Derive from Distinct Hematopoietic Lineages. Cell Reports, 3(5), 1539–1552. 10.1016/J.CELREP.2013.04.025

22. Gratwohl, A., Pasquini, M. C., Aljurf, M., Atsuta, Y., Baldomero, H., Foeken, L., Gratwohl, M., Bouzas, L. F., Confer, D., Frauendorfer, K., Gluckman, E., Greinix, H., Horowitz, M., Iida, M., Lipton, J., Madrigal, A., Mohty, M., Noel, L., Novitzky, N., … Niederwieser, D. (2015). One million haemopoietic stem-cell transplants: a retrospective observational study. The Lancet Haematology, 2(3), e91–e100. 10.1016/S2352-3026(15)00028-9

23. Hannum, G., Guinney, J., Zhao, L., Zhang, L., Hughes, G., Sadda, S. V., Klotzle, B., Bibikova, M., Fan, J. B., Gao, Y., Deconde, R., Chen, M., Rajapakse, I., Friend, S., Ideker, T., & Zhang, K. (2013). Genome-wide Methylation Profiles Reveal Quantitative Views of Human Aging Rates. Molecular Cell, 49(2), 359–367. 10.1016/J.MOLCEL.2012.10.016

24. Hemann, M. T., Strong, M. A., Hao, L. Y., & Greider, C. W. (2001). The Shortest Telomere, Not Average Telomere Length, Is Critical for Cell Viability and Chromosome Stability. Cell, 107(1), 67–77. 10.1016/S0092-8674(01)00504-9

25. Hess, N. J., Lindner, P. N., Vazquez, J., Grindel, S., Hudson, A. W., Stanic, A. K., Ikeda, A., Hematti, P., & Gumperz, J. E. (2020). Different Human Immune Lineage Compositions Are Generated in Non-Conditioned NBSGW Mice Depending on HSPC Source. Frontiers in Immunology, 11, 573406. 10.3389/FIMMU.2020.573406/TEXT

26. Higgins-Chen, A. T., Thrush, K. L., Wang, Y., Minteer, C. J., Kuo, P. L., Wang, M., Niimi, P., Sturm, G., Lin, J., Moore, A. Z., Bandinelli, S., Vinkers, C. H., Vermetten, E., Rutten, B. P. F., Geuze, E., Okhuijsen-Pfeifer, C., van der Horst, M. Z., Schreiter, S., Gutwinski, S., … Levine, M. E. (2022). A computational solution for bolstering reliability of epigenetic clocks: implications for clinical trials and longitudinal tracking. Nature Aging 2022 2:7, 2(7), 644–661. 10.1038/s43587-022-00248-2

27. Horvath, H., & Horvath, S. (2013). DNA methylation age of human tissues and cell types. Genome Biology 2013 14:10, 14(10), 3156-. 10.1186/GB-2013-14-10-R115

28. Hoshino, A., Horvath, S., Sridhar, A., Chitsazan, A., & Reh, T. A. (2019). Synchrony and asynchrony between an epigenetic clock and developmental timing. Scientific Reports 2019 9:1, 9(1), 3770-. 10.1038/s41598-019-39919-3

29. Imm, J., Pishva, E., Ali, M., Kerrigan, T. L., Jeffries, A., Burrage, J., Glaab, E., Cope, E. L., Jones, K. M., Allen, N. D., & Lunnon, K. (2021). Characterization of DNA Methylomic Signatures in Induced Pluripotent Stem Cells During Neuronal Differentiation. Frontiers in Cell and Developmental Biology, 9, 647981. 10.3389/FCELL.2021.647981/TEXT

30. Ivanovs, A., Rybtsov, S., Welch, L., Anderson, R. A., Turner, M. L., & Medvinsky, A. (2011). Highly potent human hematopoietic stem cells first emerge in the intraembryonic aorta-gonad-mesonephros region. Journal of Experimental Medicine, 208(12), 2417–2427. 10.1084/JEM.20111688

31. Kabacik, S., Lowe, D., Fransen, L., Leonard, M., Ang, S. L., Whiteman, C., Corsi, S., Cohen, H., Felton, S., Bali, R., Horvath, S., & Raj, K. (2022). The relationship between epigenetic age and the hallmarks of aging in human cells. Nature Aging 2022 2:6, 2(6), 484–493. 10.1038/s43587-022-00220-0

32. Kaur, D., Lee, S. M., Goldberg, D., Spix, N. J., Hinoue, T., Li, H.-T., Dwaraka, V. B., Smith, R., Shen, H., Liang, G., Renke, N., Laird, P. W., & Zhou, W. (2023). Comprehensive evaluation of the Infinium human MethylationEPIC v2 BeadChip. Epigenetics Communications 2023 3:1, 3(1), 6-. 10.1186/S43682-023-00021-5

33. Kollman, C., Spellman, S. R., Zhang, M. J., Hassebroek, A., Anasetti, C., Antin, J. H., Champlin, R. E., Confer, D. L., DiPersio, J. F., Fernandez-Viña, M., Hartzman, R. J., Horowitz, M. M., Hurley, C. K., Karanes, C., Maiers, M., Mueller, C. R., Perales, M. A., Setterholm, M., Woolfrey, A. E., … Eapen, M. (2016). The effect of donor characteristics on survival after unrelated donor transplantation for hematologic malignancy. Blood, 127(2), 260–267. 10.1182/BLOOD-2015-08-663823

34. Kuranda, K., Vargaftig, J., de la Rochere, P., Dosquet, C., Charron, D., Bardin, F., Tonnelle, C., Bonnet, D., & Goodhardt, M. (2011). Age-related changes in human hematopoietic stem/progenitor cells. Aging Cell, 10(3), 542–546. 10.1111/J.1474-9726.2011.00675.X;PAGEGROUP:STRING:PUBLICATION

35. Levine, M. E., Lu, A. T., Quach, A., Chen, B. H., Assimes, T. L., Bandinelli, S., Hou, L., Baccarelli, A. A., Stewart, J. D., Li, Y., Whitsel, E. A., Wilson, J. G., Reiner1, A. P., Aviv1, A., Lohman, K., Liu, Y., Ferrucci, L., & Horvath, S. (2018). An epigenetic biomarker of aging for lifespan and healthspan. Aging, 10(4), 573–591. 10.18632/AGING.101414

36. Lin, A., Giosan, I., Aparicio, A., Guo, T., Melnikas, M., Balagué-Dobón, L., Carreras-Gallo, N., Hassouneh, S. A. D., Seale, K., Kowalewski, A., Harrison, B., Smith, R., Lima Camillo, L. P. de, Lasky-Su, J., & Dwaraka, V. B. (2026). DeepStrataAge: an interpretable deep-learning clock that reveals stage- and sex-divergent DNA methylation aging dynamics. Npj Aging 2026 12:1, 12(1), 62-. 10.1038/s41514-026-00358-w

37. Luo, Q., Dwaraka, V. B., Chen, Q., Tong, H., Zhu, T., Seale, K., Raffaele, J. M., Zheng, S. C., Mendez, T. L., Chen, Y., Carreras, N., Begum, S., Mendez, K., Voisin, S., Eynon, N., Lasky-Su, J. A., Smith, R., & Teschendorff, A. E. (2023). A meta-analysis of immune-cell fractions at high resolution reveals novel associations with common phenotypes and health outcomes. Genome Medicine 2023 15:1, 15(1), 59-. 10.1186/S13073-023-01211-5

38. Marion, R. M., Strati, K., Li, H., Tejera, A., Schoeftner, S., Ortega, S., Serrano, M., & Blasco, M. A. (2009). Telomeres Acquire Embryonic Stem Cell Characteristics in Induced Pluripotent Stem Cells. Cell Stem Cell, 4(2), 141–154. 10.1016/J.STEM.2008.12.010

39. McIntosh, B. E., Brown, M. E., Duffin, B. M., Maufort, J. P., Vereide, D. T., Slukvin, I. I., & Thomson, J. A. (2015). Nonirradiated NOD,B6.SCID Il2rγ−/− KitW41/W41 (NBSGW) Mice Support Multilineage Engraftment of Human Hematopoietic Cells. Stem Cell Reports, 4(2), 171–180. 10.1016/J.STEMCR.2014.12.005

40. Mendez, E. F., Grimm, S. L., Stertz, L., Gorski, D., Movva, S. v., Najera, K., Moriel, K., Meyer, T. D., Fries, G. R., Coarfa, C., & Walss-Bass, C. (2023). A human stem cell-derived neuronal model of morphine exposure reflects brain dysregulation in opioid use disorder: Transcriptomic and epigenetic characterization of postmortem-derived iPSC neurons. Frontiers in Psychiatry, 14, 1070556. 10.3389/FPSYT.2023.1070556/TEXT

41. Ng, E. S., Sarila, G., Li, J. Y., Edirisinghe, H. S., Saxena, R., Sun, S., Bruveris, F. F., Labonne, T., Sleebs, N., Maytum, A., Yow, R. Y., Inguanti, C., Motazedian, A., Calvanese, V., Capellera-Garcia, S., Ma, F., Nim, H. T., Ramialison, M., Bonifer, C., … Elefanty, A. G. (2024). Long-term engrafting multilineage hematopoietic cells differentiated from human induced pluripotent stem cells. Nature Biotechnology 2024 43:8, 43(8), 1274–1287. 10.1038/s41587-024-02360-7

42. Onizuka, M., Imanishi, T., Harada, K., Aoyama, Y., Amaki, J., Toyosaki, M., Machida, S., Kikkawa, E., Yamada, S., Nakabayashi, K., Hata, K., Higashimoto, K., Soejima, H., & Ando, K. (2023). Donor cord blood aging accelerates in recipients after transplantation. Scientific Reports 2023 13:1, 13(1), 2603-. 10.1038/s41598-023-29912-2

43. Pang, W. W., Price, E. A., Sahoo, D., Beerman, I., Maloney, W. J., Rossi, D. J., Schrier, S. L., & Weissman, I. L. (2011). Human bone marrow hematopoietic stem cells are increased in frequency and myeloid-biased with age. Proceedings of the National Academy of Sciences of the United States of America, 108(50), 20012–20017. 10.1073/PNAS.1116110108;WEBSITE:WEBSITE:PNAS-SITE;ISSUE:ISSUE:DOI

44. Ross, J. B., Myers, L. M., Noh, J. J., Collins, M. M., Carmody, A. B., Messer, R. J., Dhuey, E., Hasenkrug, K. J., & Weissman, I. L. (2024). Depleting myeloid-biased haematopoietic stem cells rejuvenates aged immunity. Nature 2024 628:8006, 628(8006), 162–170. 10.1038/s41586-024-07238-x

45. Rossi, D. J., Bryder, D., Zahn, J. M., Ahlenius, H., Sonu, R., Wagers, A. J., & Weissman, I. L. (2005). Cell intrinsic alterations underlie hematopoietic stem cell aging. Proceedings of the National Academy of Sciences of the United States of America, 102(26), 9194–9199. 10.1073/PNAS.0503280102;WGROUP:STRING:PUBLICATION

46. Rübe, C. E., Fricke, A., Widmann, T. A., Fürst, T., Madry, H., Pfreundschuh, M., & Rübe, C. (2011). Accumulation of DNA Damage in Hematopoietic Stem and Progenitor Cells during Human Aging. PLOS ONE, 6(3), e17487. 10.1371/JOURNAL.PONE.0017487

47. Sehgal, R., Markov, Y., Qin, C., Meer, M., Hadley, C., Shadyab, A. H., Casanova, R., Manson, J. A. E., Bhatti, P., Moore, A. Z., Crimmins, E. M., Hagg, S., Assimes, T. L., Whitsel, E. A., Higgins-Chen, A. T., & Levine, M. (2025). Systems Age: a single blood methylation test to quantify aging heterogeneity across 11 physiological systems. Nature Aging 2025 5:9, 5(9), 1880–1896. 10.1038/s43587-025-00958-3

48. Shaw, B. E., Logan, B. R., Spellman, S. R., Marsh, S. G. E., Robinson, J., Pidala, J., Hurley, C., Barker, J., Maiers, M., Dehn, J., Wang, H., Haagenson, M., Porter, D., Petersdorf, E. W., Woolfrey, A., Horowitz, M. M., Verneris, M., Hsu, K. C., Fleischhauer, K., & Lee, S. J. (2018). Development of an Unrelated Donor Selection Score Predictive of Survival after HCT: Donor Age Matters Most. Biology of Blood and Marrow Transplantation, 24(5), 1049–1056. 10.1016/j.bbmt.2018.02.006

49. Snowden, J. A., Sánchez-Ortega, I., Corbacioglu, S., Basak, G. W., Chabannon, C., de la Camara, R., Dolstra, H., Duarte, R. F., Glass, B., Greco, R., Lankester, A. C., Mohty, M., Neven, B., de Latour, R. P., Pedrazzoli, P., Peric, Z., Yakoub-Agha, I., Sureda, A., & Kröger, N. (2022). Indications for haematopoietic cell transplantation for haematological diseases, solid tumours and immune disorders: current practice in Europe, 2022. Bone Marrow Transplantation 2022 57:8, 57(8), 1217–1239. 10.1038/s41409-022-01691-w

50. Søraas, A., Matsuyama, M., de Lima, M., Wald, D., Buechner, J., Gedde-Dahl, T., Søraas, C. L., Chen, B., Ferrucci, L., Dahl, J. A., Horvath, S., & Matsuyama, S. (2019). Epigenetic age is a cell-intrinsic property in transplanted human hematopoietic cells. Aging Cell, 18(2), e12897. 10.1111/ACEL.12897;JOURNAL:JOURNAL:14749726;PAGE:STRING:ARTICLE/CHAPTER

51. Steg, L. C., Shireby, G. L., Imm, J., Davies, J. P., Franklin, A., Flynn, R., Namboori, S. C., Bhinge, A., Jeffries, A. R., Burrage, J., Neilson, G. W. A., Walker, E. M., Perfect, L. W., Price, J., McAlonan, G., Srivastava, D. P., Bray, N. J., Cope, E. L., Jones, K. M., … Hannon, E. (2021). Novel epigenetic clock for fetal brain development predicts prenatal age for cellular stem cell models and derived neurons. Molecular Brain 2021 14:1, 14(1), 98-. 10.1186/S13041-021-00810-W

52. Stölzel, F., Brosch, M., Horvath, S., Kramer, M., Thiede, C., von Bonin, M., Ammerpohl, O., Middeke, M., Schetelig, J., Ehninger, G., Hampe, J., & Bornhäuser, M. (2017). Dynamics of epigenetic age following hematopoietic stem cell transplantation. Haematologica, 102(8), e321–e323. 10.3324/HAEMATOL.2016.160481

53. Suhr, S. T., Chang, E. A., Rodriguez, R. M., Wang, K., Ross, P. J., Beyhan, Z., Murthy, S., & Cibelli, J. B. (2009). Telomere Dynamics in Human Cells Reprogrammed to Pluripotency. PLOS ONE, 4(12), e8124. 10.1371/JOURNAL.PONE.0008124

54. Sun, D., Luo, M., Jeong, M., Rodriguez, B., Xia, Z., Hannah, R., Wang, H., Le, T., Faull, K. F., Chen, R., Gu, H., Bock, C., Meissner, A., Göttgens, B., Darlington, G. J., Li, W., & Goodell, M. A. (2014). Epigenomic Profiling of Young and Aged HSCs Reveals Concerted Changes during Aging that Reinforce Self-Renewal. Cell Stem Cell, 14(5), 673–688. 10.1016/J.STEM.2014.03.002

55. Takahashi, K., Tanabe, K., Ohnuki, M., Narita, M., Ichisaka, T., Tomoda, K., & Yamanaka, S. (2007). Induction of Pluripotent Stem Cells from Adult Human Fibroblasts by Defined Factors. Cell, 131(5), 861–872. 10.1016/J.CELL.2007.11.019

56. Vaziri, H., Dragowska, W., Allsopp, R. C., Thomas, T. E., Harley, C. B., & Lansdorp, P. M. (1994). Evidence for a mitotic clock in human hematopoietic stem cells: Loss of telomeric DNA with age. Proceedings of the National Academy of Sciences of the United States of America, 91(21), 9857–9860. 10.1073/PNAS.91.21.9857;PAGE:STRING:ARTICLE/CHAPTER

57. Vaziri, H., Schächter, F., Uchida, I., Wei, L., Zhu, X., Effros, R., Cohen, D., & Harley, C. B. (1993). Loss of telomeric DNA during aging of normal and trisomy 21 human lymphocytes. American Journal of Human Genetics, 52(4), 661. https://pmc.ncbi.nlm.nih.gov/articles/PMC1682068/

58. Vodyanik, M. A., Bork, J. A., Thomson, J. A., & Slukvin, I. I. (2005). Human embryonic stem cell–derived CD34+ cells: efficient production in the coculture with OP9 stromal cells and analysis of lymphohematopoietic potential. Blood, 105(2), 617–626. 10.1182/BLOOD-2004-04-1649

59. Weidner, C. I., Lin, Q., Koch, C. M., Eisele, L., Beier, F., Ziegler, P., Bauerschlag, D. O., Jöckel, K. H., Erbel, R., Mühleisen, T. W., Zenke, M., Brümmendorf, T. H., & Wagner, W. (2014). Aging of blood can be tracked by DNA methylation changes at just three CpG sites. Genome Biology 2014 15:2, 15(2), R24-. 10.1186/GB-2014-15-2-R24

60. Weidner, C. I., Ziegler, P., Hahn, M., Brümmendorf, T. H., Ho, A. D., Dreger, P., & Wagner, W. (2014). Epigenetic aging upon allogeneic transplantation: the hematopoietic niche does not affect age-associated DNA methylation. Leukemia 2015 29:4, 29(4), 985–988. 10.1038/leu.2014.323

61. Yamaguchi, H., Calado, R. T., Ly, H., Kajigaya, S., Baerlocher, G. M., Chanock, S. J., Lansdorp, P. M., & Young, N. S. (2005). Mutations in TERT, the Gene for Telomerase Reverse Transcriptase, in Aplastic Anemia. New England Journal of Medicine, 352(14), 1413–1424. 10.1056/NEJMOA042980;REQUESTEDJOURNAL:JOURNAL:NEJM;PAGE:STRING:ARTICLE/CHAPTER

62. Yanai, H., McNeely, T., Ayyar, S., Leone, M., Zong, L., Park, B., & Beerman, I. (2024). DNA methylation drives hematopoietic stem cell aging phenotypes after proliferative stress. GeroScience 2024 47:2, 47(2), 1873–1886. 10.1007/S11357-024-01360-4

63. Yin, A. H., Miraglia, S., Zanjani, E. D., Almeida-Porada, G., Ogawa, M., Leary, A. G., Olweus, J., Kearney, J., & Buck, D. W. (1997). AC133, a Novel Marker for Human Hematopoietic Stem and Progenitor Cells. Blood, 90(12), 5002–5012. 10.1182/BLOOD.V90.12.5002

64. Zbieć-Piekarska, R., Spólnicka, M., Kupiec, T., Makowska, Z., Spas, A., Parys-Proszek, A., Kucharczyk, K., Płoski, R., & Branicki, W. (2015). Examination of DNA methylation status of the ELOVL2 marker may be useful for human age prediction in forensic science. Forensic Science International: Genetics, 14, 161–167. 10.1016/j.fsigen.2014.10.002

65. Zeng, Y., He, J., Bai, Z., Li, Z., Gong, Y., Liu, C., Ni, Y., Du, J., Ma, C., Bian, L., Lan, Y., & Liu, B. (2019). Tracing the first hematopoietic stem cell generation in human embryo by single-cell RNA sequencing. Cell Research 2019 29:11, 29(11), 881–894. 10.1038/s41422-019-0228-6

